# Influenza virus infection in the lungs leads to pancytopenia and defective immune cell differentiation program in the thymus and bone marrow

**DOI:** 10.1101/2025.04.22.650071

**Authors:** Prajakta Shinde, Giovannino Silvestri, Panjamurthy Kuppusamy, Nicholas Stamatos, Chozha Vendan Rathinam

## Abstract

Exaggerated inflammation and cytokine storm are hallmark features of influenza A virus (IAV)-induced respiratory diseases. While previous studies unequivocally demonstrated the pathophysiological consequences (multiorgan failure) of IAV-associated cytokine storm, it remains unknown if IAV-induced systemic inflammation impacts the “fitness” and differentiation of immune cells from hematopoietic stem cells (HSCs). Our data on lethal IAV-infected C57BL/6 wildtype mice after 10 days of infection indicated reduced monocyte- and lymphocyte-counts in the peripheral blood, and overall cellularity of spleen, thymus and lymph nodes. IAV-infection resulted in increased numbers of myeloid cells, CD8^+^ T cells, alveolar macrophages (AVMs), CD11b^+^ dendritic cells (DCs) & plasmacytoid DCs (pDCs), whereas decreased frequencies of CD103^+^ DCs, in the lungs of IAV-infected mice. Analysis of spleen and draining lymph nodes indicated reduced absolute numbers of B cells, T cells, monocytes and DCs after 10 days of lethal IAV infection. Thymic analysis indicated perturbed T cell differentiation and bone marrow (BM) data revealed impaired DC differentiation following IAV infection. Hematopoietic stem and progenitor cells (HSPCs) studies demonstrated an imbalanced distribution of HSCs, multipotent progenitors (MPPs), myeloid progenitors and DC progenitors within the BM niche. Mechanistic studies exhibited elevated levels of systemic inflammation and altered local ‘pro-inflammatory milieu’. Molecular analyses documented elevated levels of intracellular reactive oxygen species (ROS) at all stages of HSPC differentiation and increased mass of active mitochondria in HSPC subsets. In essence, our studies provide novel insights into mechanisms through which lethal IAV-infection induces deficiencies of the innate and adaptive immune system.

## Introduction

Influenza A virus (IAV) causes severe pathology that mainly affects infants, pregnant women, and the elderly or immunocompromised individuals(1, 2). The exaggerated inflammatory responses, produced by a variety of hos cells, such as epithelial cells, endothelial cells, and immune cells, during an acute phase of IAV infection have been termed as the “cytokine storm”. Even though this augmented inflammation serves critical functions at the acute phases of infection and helps the clearance of IAV, cytokine storm often results in widespread tissue damage(3). More importantly, cytokines storm is believed to be one of the key contributing factors for increased IAV infection-associated mortalities(4-7), major immunopathology, systemic inflammation and serious disease consequences(5, 8-10), including multiple organ dysfunction syndromes(11-13).

During IAV infection, the first wave of immune responses and efficient recruitment of immune cells in the airways are created by alveolar macrophages (AVM)(14). This is followed by the influx of neutrophils and secretion of inflammatory mediators by resident immune cells, such as dendritic cells (DCs), monocytes and macrophages, in the lungs, which serve as a fundamental defense mechanism(15-17). Control of infection and clearance of IAV depend on pathogen specific adaptive immune responses(18). While cytotoxic T-lymphocyte (CTL) activity has been shown to be both necessary and sufficient for recovery from influenza(19), involvement of both CD4^+^ and CD8^+^ T cells are needed for the immunity against IAV. In view of the fact that the strength and specificity of T (CD8 and CD4)-cell mediated immunity is dependent on efficient presentation of IAV-associated antigens by DCs (and macrophages)(20, 21), functions of innate myeloid cells are believed to the key in determining the outcome of IAV infection(1, 22-26). Indeed, disease severity in hospitalized patients has been shown to correlate with monocytes recruitment with increased levels of cytokines(2), strong TNF-producing monocytic responses in the blood(1) and inflammatory, neutrophil-dominant patterns(27). Furthermore, DCs and monocytes are recruited to the nasopharynx of individuals hospitalized with severe influenza infections and infected with 2009 H1N1pdm IAV strains(2, 28, 29). These studies highlight a vital role and an involvement of innate myeloid cells in the immunity against IAV infection.

Here, we demonstrate that IAV infection affects the composition of innate myeloid cells in the lungs and alters distribution and numbers of both innate and adaptive immune subsets in the periphery. Interestingly, IAV infection causes a profound impact on the differentiation and maintenance of the DC-lineage cells and alters the immunogenicity of all DC subsets in the secondary and peripheral lymphoid organs. Our data specify that maintenance of hematopoietic stem and progenitor cells (HSPCs) and their differentiation into DC progenitors were altered in during IAV infection. Mechanistic studies suggested increased systemic and local inflammation, augmented mitochondrial mass and elevated levels of intracellular (reactive oxygen species) ROS in HSPCs and immune cells. Overall, our studies demonstrate that IAV infection interferes with early hematopoietic and immune differentiation program in the bone marrow (BM) and thymus.

## Materials and Methods

### Mice

C57BL6/J mice were purchased from the Jackson Laboratory. The Institutional Animal Care and Use Committee approved all mouse experiments.

### Influenza virus

Influenza A/ Netherlands/602/2009 (NL09; H1N1) virus was a kind gift from Dr Matthew Frieman (University of Maryland School of Medicine). The virus was grown in Madine Darby canine kidney cells as per standard protocol and was purified, titered, and stored at −80°C.

### Mouse infection with Influenza virus

All experiments were performed on 6 - 8-week-old, age- and sex-matched C57BL/6J mice (Jackson Laboratories, USA) that were handled in accordance with protocols approved by the Institutional Animal Care and Use Committee (IACUC) of the University of Maryland School of Medicine. Prior to inoculation, frozen stocks of NL09 were diluted to 100 PFU/ml in sterile, endotoxin-free, phosphate buffered saline (PBS, Biosource International, Rockville, MD). Mice were lightly anesthetized with 4% isoflurane (VETone, Boise, ID) and 2% 02 delivered by a regulated isoflurane vaporizer (VETequip, Livermore, CA) and 10 μl of inoculum (for a total dose of 1000 PFU) was administered to a single nares of each mouse, as described previously (30). Mice inoculated with PBS alone were used as controls. Each mouse was observed daily for weight change and clinical signs of moribundity or euthanized to harvest tissue for further analysis as per IACUC guidelines.

### Blood collection & analysis

Peripheral blood from mice was collected through sub-mandibular puncture in tubes containing EDTA. Complete blood count analysis was performed using Element HT5 Veterinary Hematology Analyzer (Heska).

### Single cell preparation

Organs from IAV-infected and uninfected mice were harvested after 10 days of infection. Spleen, thymus and Lungs were cut into small pieces and digested with 0.5 mg/mL Collagenase D + 50 U/mL DNase I for 45 mins at 37°C. Single cell preparation was made after mincing and passing through the cell strainer using ice cold PBS with 2% FCS. RBCs were lysed with ammonium chloride (STEMCELL Technologies). Trypan blue (Amresco)–negative cells were counted as live cells.

### Flow cytometry

Cells were analyzed by flow cytometry with Attune NxT (Thermofisher) and FlowJo software (Tree Star). The following monoclonal antibodies were used: anti-CD34 (RAM34), anti-CD48 (HM48-1), anti-CD117 (2B8), anti-Flt3 (A2F10.1), anti-Sca-1 (D7), anti-B220 (RA3-6B2), anti-CD19 (1D3), anti-CD3 (145-2C11), anti-CD4 (GK1.5), anti-CD8 (53-6.7), anti-CD11b (M1/70), anti– Gr-1 (RB6-8C5) and anti-Ter119 (TER119; from BD Biosciences); anti-CD150 (TC15-12F12.2) from Biolegend; anti-CD16/32 (93) from eBioscience; anti-CD11c (N418), anti-CD8(53-6.7), anti-PDCA1 (129C1), anti-CD24 (M1/69), anti-SIRPa (P84), anti-CD103 (2E7), anti-CD45 (S18009F), anti-CD207 (4C7), anti-CD115 (AFS98), anti-LY6C (HK1.4), anti-SiglecH (551), anti-CLEC9A (7H11), anti-CD80 (16-10A1), anti-CD86 (GL-1), anti-CD40(3/23), anti-H2K^b^ (AF6-88.5), anti-IA/IE (M5/114.15.2), anti-CD26 (H194-112), anti-XCR1 (Zet) from Biolegend. Cells incubated with biotinylated monoclonal antibodies were incubated with fluorochrome-conjugated streptavidin–peridinin chlorophyll protein–cyanine 5.5 (551419; BD) or streptavidin-allophycocyanin-Cy7 (554063; BD). In all the FACS plots, indicated are the percentages (%) of the gated fraction.

### RNA extraction, PCR, and real-time PCR

Total RNA was isolated with a RNeasy Mini kit or RNeasy Micro kit (QIAGEN). cDNA was synthesized with Oligo(dT) primer and Maxima reverse transcription (Thermo Fisher Scientific). PCR was performed with T100 thermal cycler (Bio-Rad Laboratories) and TSG Taq (Lamda Biotech). Real-time PCR was performed in duplicates using gene specific primers with a CFX-connect real-time PCR system (Bio-Rad Laboratories) and iTaq SYBR Green Supermix (Bio-Rad Laboratories) according to the manufacturer’s instructions. Relative expression was normalized to the expression levels of the internal control (housekeeping gene) Hprt.

### ELISA studies

Levels of inflammatory cytokines in the blood sera were analyzed using ELISA kits (BD Biosciences), according to manufacturer’s instructions.

### Measuring ROS levels and mitochondrial mass

Cells were stained with cell surface markers and then incubated with 2mM CM-H_2_DCFDA (Life Technologies C6827) and MitoTracker Green FM (Life Technologies M46750) in pre-warmed HBSS at 37°C for 15-20 min. The cells were then pelleted and resuspended in PBS before acquisition.

### Statistics

Data represent mean and s.e.m Two-way ANOVA, Log RANK test, and Two-tailed student’s t tests were used to assess statistical significance (n.s. = not significant, *P < 0.05, **P<0.01, *** P< 0.001 and **** P< 0.0001).

## Results

### IAV infection alters the distribution and functions of innate myeloid cells within the lungs

To study IAV-induced hematopoietic and immune defects, C57BL/6 wildtype mice were infected with IAV on Day 0 and analyzed thereafter. IAV-infected mice exhibited significant weight loss starting at 3 days of infection (**Data not shown**) and ∼ 50 % of the mice died within 7 days of infection and almost all mice died by 15 days of infection (**Fig. 1, A**). To study and understand IAV-induced pathophysiological impact on the immune system, we analyzed mice that survived between 8 & 10 days of IAV infection. Complete blood count (CBC) analysis revealed a reduction in total white blood cells (WBC) counts, particularly lymphocytes, monocytes and eosinophils (**Fig. 1, B & C**), and an increase of platelets, red blood cells (RBCs), hemoglobin and hematocrit levels (**Fig. 1, D**) in IAV-infected mice. Consistently, Flow cytometric analysis indicated an increase of CD11b^+^Ly6G^+^ granulocytes, and a decrease of CD11c^+^MHC-CII^+^ DCs, CD19^+^ B cells, CD4^+^ T cells and CD8^+^ T cells in the peripheral blood of IAV-infected mice (**Fig. S1, A**). Enumeration of total cell counts of primary and secondary lymphoid organs from IAV-infected and uninfected mice, after 10 days of infection indicated modestly reduced cellularity of the BM (though not statistically significant), but strikingly reduced cellularity of the spleen, thymus, and peripheral lymph nodes of IAV-infected mice (**Fig. 1,E**).

**Figure 1.**
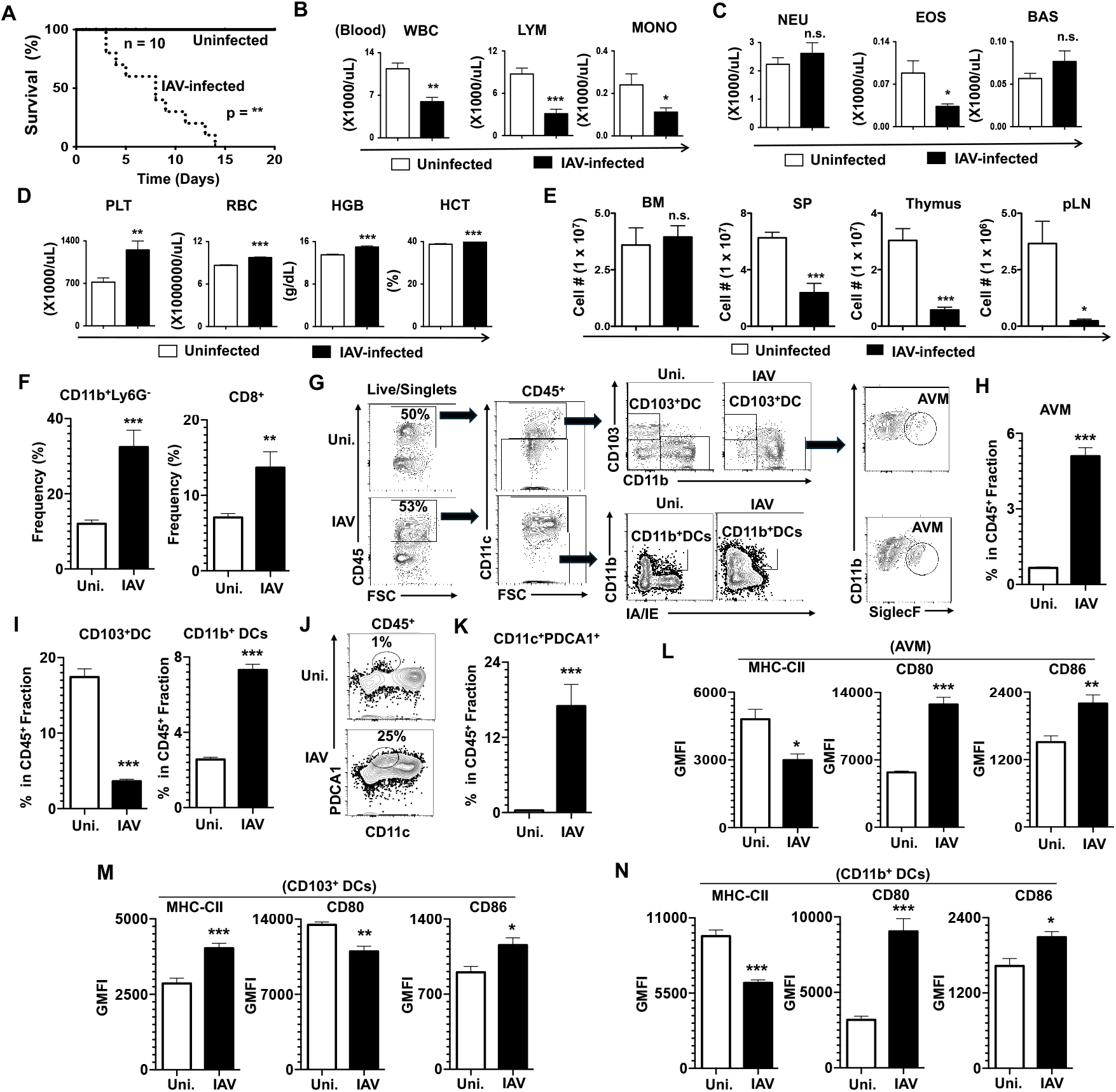
IAV infection results in pancytopenia and altered immune distribution in lungs. **A**. Kaplan and Meier survival curve analysis of uninfected and IAV-infected mice (n= 10). **B**-**D**. Complete blood count analysis of peripheral blood from uninfected and IAV-infected mice (n=6). **E**. Absolute cell counts of BM, spleen, thymus and peripheral lymph nodes (pLN) from uninfected and IAV-infected mice (n= 6). **F**. Frequencies of indicated immune subsets in the lungs of uninfected and IAV-infected mice (n= 6). **G**. FACS plots indicating the gating strategy of identifying CD103^+^DCs, CD11b^+^DCs and AVM within the lungs of uninfected and IAV-infected mice (n= 6). **H & I**. Frequencies of AVM (**H**), CD103^+^DCs (**I, left**) & CD11b^+^DCs (**I, right**) within CD45^+^ fraction of the lungs from uninfected and IAV-infected mice (n=6). **J & K**. FACS plots indicating gating strategy for identifying pDCs (**J**) and Frequencies (**K**) within CD45^+^ fraction of pDCs in the lungs of uninfected and IAV-infected mice (n=6). **L-N**. Surface expression levels of MHC-CII, CD80 and CD86 in total AVM (**L**), CD103^+^DCs (**M**) and CD11b^+^DCs (**N**) from the lungs of uninfected and IAV-infected mice (n=6). Shown were Geo-Mean Fluorescence Intensities (GMFI). Shown are data from mice after 10 days of IAV infection (**C-N**). All data represent mean ± SEM. Two-way ANOVA (**A**), Log RANK test (**B**), and Two-tailed student’s t tests (**C-N**) were used to assess statistical significance (n. s. = not significant, *P < 0.05, **P<0.01, *** P< 0.001 & **** P< 0.0001).

To identify immune alterations in the lungs following lethal IAV infection, we performed flow cytometric analysis. Our immunophenotyping studies of the lungs indicated normal frequencies of CD11b^+^Ly6G^+^ granulocytes, CD19^+^ B cells and CD4^+^ T cells (**Fig. S1, A**), but remarkably increased frequencies of CD8^+^ T cells and CD11b^+^Ly6G^-^ myeloid cells (**Fig. 1F**), within the total CD45^+^ fraction of the lungs of IAV-infected mice. Next, we determined the frequencies of CD45^+^CD11c^high^CD103^-^ SiglecF^+^CD11b^+^alveolar macrophages (AVMs) and our data indicated a striking increase of AVM in the lungs after 10 days of IAV infection (**Fig. 1, G & H**). Analysis of DC subsets indicated reduced frequencies of CD45^+^CD11c^+^CD11b^-^CD103^+^ DCs (equivalent of cDC1) and increased frequencies of CD45^+^CD11c^-^ CD11b^+^CII^high^ DCs (equivalent of cDC2) (**Fig. 1, G & I)**, and augmented frequencies of CD11c^int^PDCA1^+^ pDCs (**Fig. 1, J &K**), in the lungs after 10 days of IAV infection. To study alterations in immunogenicity of AVMs and DC subsets after IAV-infection, we assessed the surface expression levels of MHC-Class II, CD80 and CD86. Our data revealed; reduced expression of MHC-Class II, but increased expression of CD80 and CD86 in AVMs (**Fig. 1L**); augmented expression of MHC-Class II and CD86, but reduced expression of CD80, in CD103^+^ DCs (**Fig. 1M**); and reduced expression of MHC-Class II, but increased expression of CD80 and CD86 in CD11b^+^ DCs (**Fig. 1N**), of lungs after 10 days of IAV infection. Together, these data indicated that IAV infection causes major alterations in the innate myeloid subsets and lineage-specific alterations in frequencies and immunogenicity of CD103^+^ DCs and CD11b^+^ DCs.

### IAV infection impacts distribution and composition of immune subsets in the spleen

To determine the consequences of IAV infection on the peripheral immune system, we analyzed the alterations and composition of immune cells within the spleen. Our data indicated normal frequencies (**Fig. S2A**), but reduced absolute numbers (**Fig. 2A**), of total CD19^+^ B cells, elevated expression of IgD (**Fig. 2B**), and normal expression of IgM (**Fig. S2B**) in B cells (**Fig. 2B**) following IAV infection. T cells analysis revealed modestly increased frequencies of total T (CD90^+^TCRβ^+^) cells (**Fig. S2C**). Further analysis suggested normal frequencies (**Fig. S2D**), but reduced absolute numbers (**Fig. 2C**) of CD4 T cells, and modestly increased frequencies, but reduced absolute numbers, of CD8 T cells (**Fig. 2D**) in the spleen of IAV mice. Interestingly, analysis of CD25^+^ T cells suggested increased frequencies, but normal absolute numbers, of both CD4^+^ T (**Fig. 2E & Fig. S2E**) and CD8^+^ T (**Fig. 2F & Fig. S2F**) cells following IAV infection. Next, we enumerated the frequencies and numbers of the myeloid subsets. Immunophenotyping studies revealed normal frequencies and absolute numbers of Ly6G^+^CD11b^+^ granulocytes (**Fig. S2G**), reduced frequencies and absolute numbers of Ly6G^-^CD11b^+^ monocytes/macrophages (**Fig. 2G**) and normal frequencies and absolute numbers of Ter119^+^ erythrocytes (**Fig. S2H**) in the spleen of IAV-infected mice. Of note, frequencies and absolute numbers of CD11c^+^CII^+^ cDCs were remarkably reduced in the spleen of IAV-infected mice (**Fig. 2H & I)**. DC subset analysis indicated; increased frequencies, but reduced absolute numbers, of CD11c^+^CII^+^XCR1^+^SIRPα^-^(**Fig. 2H & J**) and CD11c^+^CII^+^CD8^+^CD11b^-^ (**Fig. 2H & L**) cDC1 subsets; and reduced frequencies and absolute numbers of CD11c^+^CII^+^XCR1^-^SIRPα^+^(**Fig. 2H & K**) and CD11c^+^CII^+^CD8^-^CD11b^+^ (**Fig. 2H & M**) cDC2 subsets in the spleen of IAV-infected mice. Analysis of CD11c^int^PDCA1^+^ pDC subset revealed strikingly reduced frequencies and absolute numbers of pDCs in the spleen of IAV-infected mice (**Fig. 2N & O**). Consistent with our previous data (**Fig. 2H-M**), our analysis of CD11c^high^PDCA1^-^ subset indicated reduced frequencies and absolute numbers of total cDCs (**Fig. 2N & Fig. S2I**) in the spleen of IAV-infected animals. Further cDC analysis indicated increased frequencies, but reduced absolute numbers, of CD11c^high^PDCA1^-^CD8^+^CD11b^-^ cDC1 subset (**Fig. S2J**), and reduced frequencies and absolute numbers of CD11c^high^PDCA1^-^CD8^-^CD11b^+^ cDC2 subset (**Fig. S2K**) in the spleen of IAV-infected mice. Next, we determined the frequencies of CD11c^high^PDCA1^-^CD8^-^CD11b^-^ precursor (pre)- cDC subset, CD11c^high^PDCA1^-^CD8^-^CD11b^-^CD24^+^SIRPα^-^ pre-cDC1 and CD11c^high^PDCA1^-^CD8^-^ CD11b^-^CD24^-^SIRPα^+^ pre-cDC2 subsets(31). Our analysis indicated; normal frequencies, but reduced absolute numbers, of pre-cDCs (**Fig. S2L**); increased frequencies, but decreased absolute numbers, of pre-cDC1 subset (**Fig. S2M**); and decreased frequencies and absolute numbers of pre-cDC2 subset (**Fig. S2N**) in the spleen of IAV-infected animals. Similarly, analysis of pDC subsets indicated reduced frequencies and absolute numbers of CD11c^low^Ly6C^+^PDCA1^-^ immature pDCs (**Fig. 2P & Q**) and increased frequencies, but reduced absolute numbers, of CD11c^low^Ly6C^+^PDCA1^+^ mature pDCs (**Fig. 2P & 2R**) in the spleen of IAV-infected mice. Finally, we determined the surface expression levels of MHC-CI, MHC-CII, CD80 and CD86 in cDC and pDC subsets. Our analysis revealed; increased expression of MHC-CI in cDC1, cDC2, and pDC subsets, albeit at different levels (**Fig. 2S top**); reduced expression of MHC-CII in cDC1 subset, but increased expression of MHC-CII in cDC2, and pDC subsets (**Fig. 2S bottom**); reduced expression of CD80 in cDC1 and cDC2 subsets, but remarkably elevated expression of CD80 in pDC subset (**Fig. 2T top**); and normal expression of CD86 in cDC1, but increased expression of CD86 in cDC2 and pDC subsets (**Fig. 2T bottom**), in the spleen of IAV-infected mice. Overall, these data indicated that IAV infection results in modest alterations of lymphoid- and myeloid-lineage subsets in the spleen and that the distribution, absolute numbers and immunogenic properties of both cDC and pDC subsets were remarkably affected by IAV infection.

**Figure 2.**
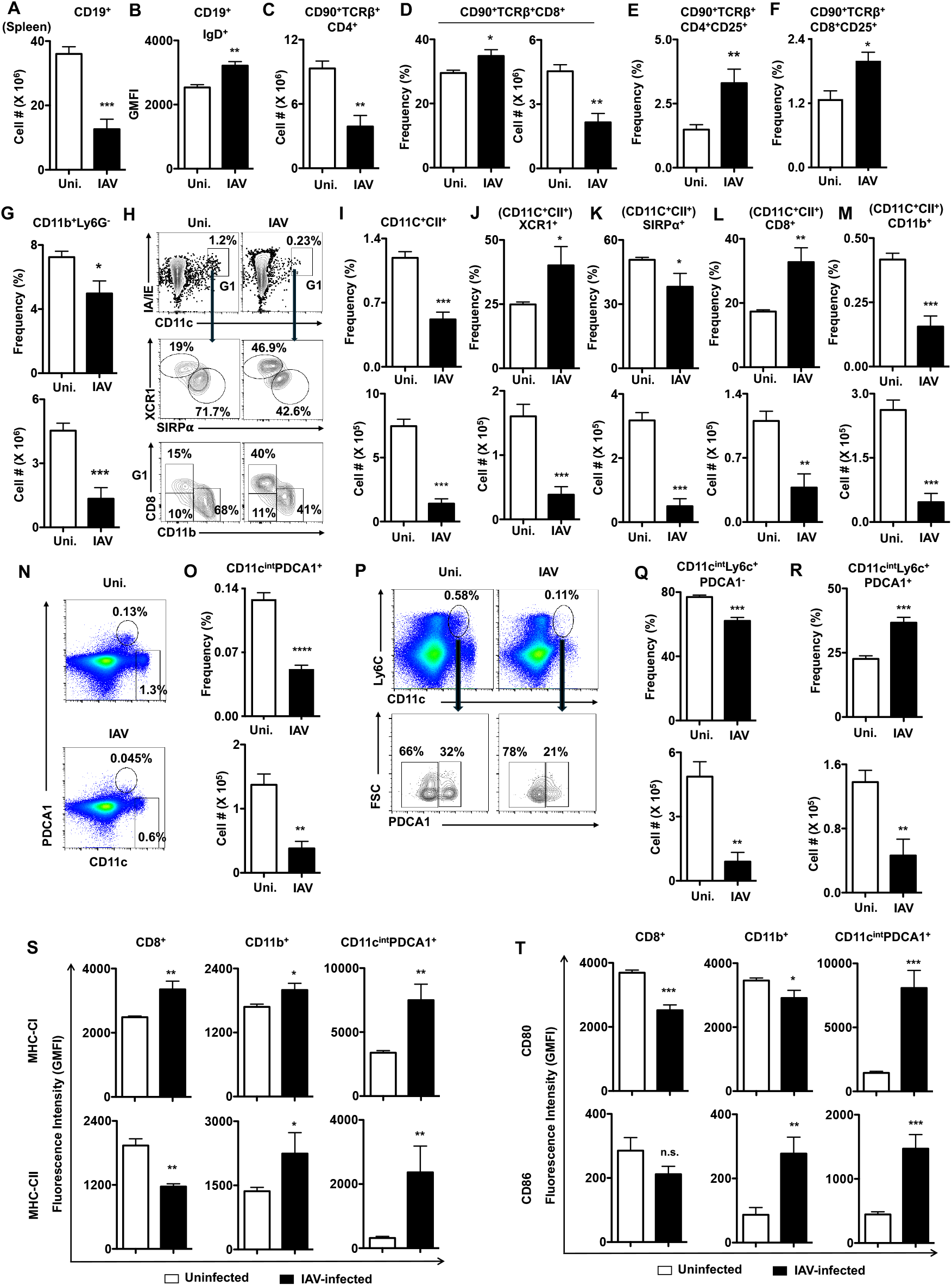
IAV infection causes alterations in DC subsets of the spleen. **A**. Absolute cell counts of CD19^+^ cells of spleen from uninfected and IAV-infected mice (n= 6). **B**. Expression levels of IgD^+^ within CD19^+^ cells of spleen from uninfected and IAV-infected mice (n= 6). **C**. Absolute cell counts of CD90^+^TCRb^+^CD4^+^ cells from the spleen from uninfected and IAV-infected mice (n= 6). **D**. Frequencies and absolute cell counts of CD90^+^TCRb^+^CD8^+^ cells from the spleen from uninfected and IAV-infected mice (n= 6). (**E & F**). Frequencies of CD90^+^TCRb^+^CD4^+^CD25^+^ cells (**E**) and CD90^+^TCRb^+^CD8^+^CD25^+^ cells (**F**) from the spleen from uninfected and IAV-infected mice (n= 6). **G**. Frequencies (top) and absolute cell counts (bottom) of CD11b^+^Ly6G^-^ cells from the spleen from uninfected and IAV-infected mice (n= 6). **H**. FACS plots indicating the gating strategy of identifying CD11c^+^CII^+^ total cDC (top), XCR1^+^ SIRPα^-^ cDC1 & XCR1^-^SIRPα^+^ cDC2 subsets (middle), and CD8^+^CD11b^-^ cDC1 & CD8^-^CD11b^+^ cDC2 subsets (bottom) within the CD11c^+^CII^+^ cells of the spleen from uninfected and IAV-infected mice (n= 6). **I-M**. Frequencies (top) and absolute cell counts (bottom) of CD11c^+^CII^+^ (**I**), CD11c^+^CII^+^XCR1^+^SIRPα^-^ (**J**), CD11c^+^CII^+^XCR1^-^SIRPα^+^ (**K**), CD11c^+^CII^+^CD8^+^CD11b^-^ (**L**) and CD11c^+^CII^+^CD8^-^CD11b^+^ (**M**) cells of the spleen from uninfected and IAV-infected mice (n= 6). **N & O**. FACS plots indicating the gating strategy of identifying CD11c^int^PDCA1^+^ pDCs (**N**), and frequencies (top) and absolute cell counts (bottom) of pDCs (**O**) from the spleen of uninfected and IAV-infected mice (n= 6). **P-R**. FACS plots indicating the gating strategy of identifying CD11c^int^ Ly6C^+^PDCA1^-^ immature pDCs and mature pDCs (**P**), and frequencies (top) and absolute cell counts (bottom) of immature pDCs (**Q**) and mature pDCs (**R**) from the spleen of uninfected and IAV-infected mice (n= 6). **S & T**. Surface expression levels (GMFI) of MHC-CI and MHC-CII (**S**), and CD80 and CD86 (**T**) in CD8^+^DCs (left), CD11b^+^DCs (middle) and CD11c^int^PDCA1^+^ pDCs from the spleen of uninfected and IAV-infected mice (n=6). Shown are data from mice after 10 days of IAV infection (**A-T**). All data represent mean ± SEM. Two-tailed student’s t tests were used to assess statistical significance (n. s. = not significant, *P < 0.05, **P<0.01, *** P< 0.001 & **** P< 0.0001).

### IAV infection affects immune functions of lymph nodes

To determine if IAV infection alters distribution of immune cells within the lymphatic system, we isolated peripheral draining lymph nodes and performed immunophenotyping studies. Our analysis indicated; normal frequencies, but reduced absolute numbers, of CD19^+^ B cells (**Fig. 3A**); reduced frequencies and absolute numbers of total CD4^+^ T cells (**Fig. 3B**); diminished absolute numbers of CD44^low/-^ CD4^+^ T cells (**Fig. 3C**); reduced frequencies and absolute numbers of total CD8^+^ T cells (**Fig. 3D**); decreased absolute numbers of CD44^low/-^ CD8^+^ T cells (**Fig. 3E**); increased frequencies, but normal absolute numbers, of CD25^+^CD4^+^ T cells (**Fig. 3F**); and increased frequencies and absolute numbers of CD25^+^CD8^+^ T cells (**Fig. 3G**) in the peripheral/draining lymph nodes of IAV-infected mice.

**Figure 3.**
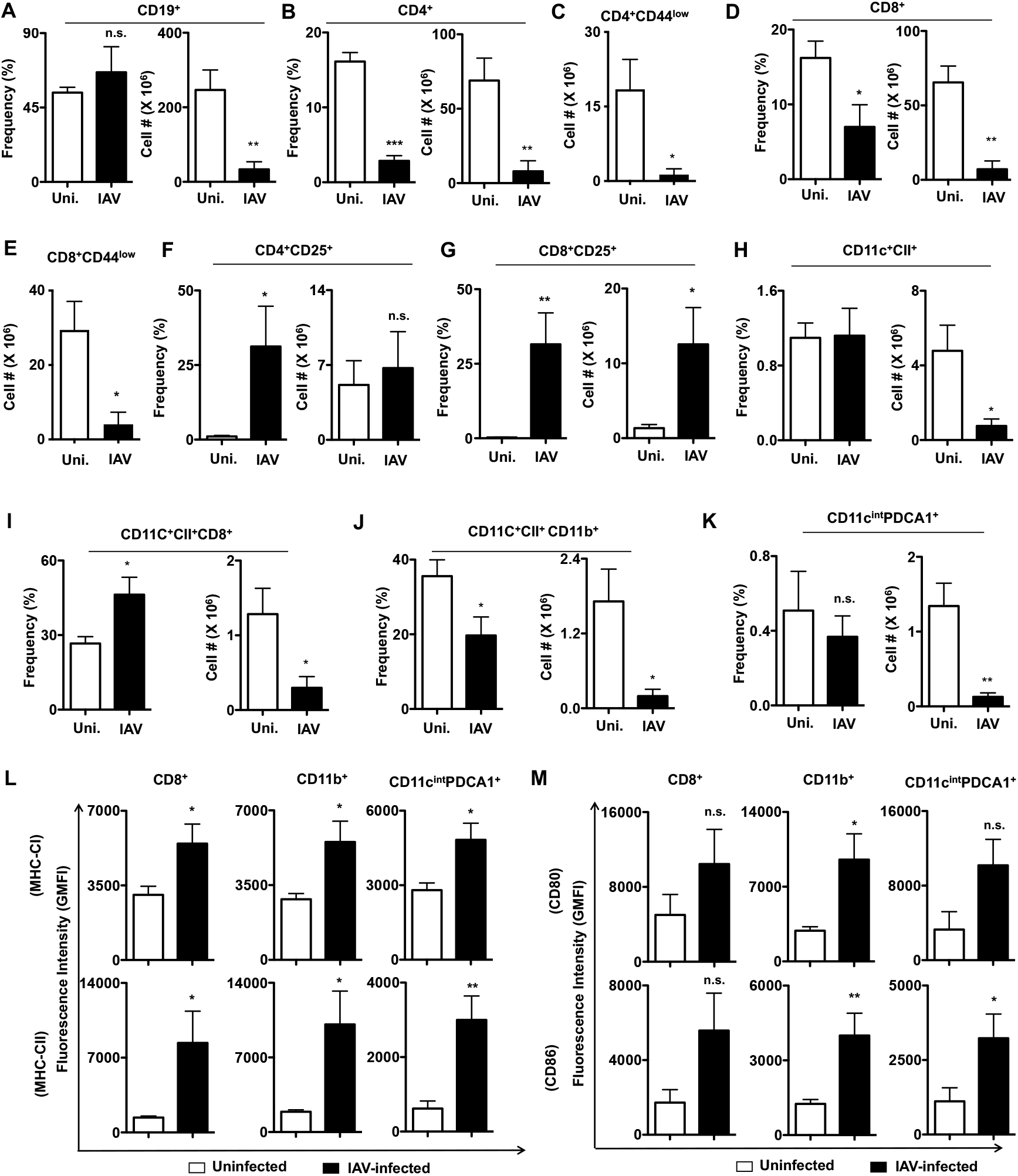
IAV infection causes immune alterations in the peripheral lymph nodes. **A & B**. Frequencies (left) and absolute cell counts (right) of CD19^+^ (**A**) and CD4^+^ (**B**) cells of pLN from uninfected and IAV-infected mice (n= 6). **C**. Absolute cell counts of CD4^+^CD44^low^ cells of pLN from uninfected and IAV-infected mice (n= 6). **D**. Frequencies (left) and absolute cell counts (right) of CD8^+^ cells of pLN from uninfected and IAV-infected mice (n= 6). **E**. Absolute cell counts of CD8^+^CD44^low^ cells of pLN from uninfected and IAV-infected mice (n= 6). **F & G**. Frequencies (left) and absolute cell counts (right) of CD4^+^CD25^+^ (**F**) and CD8^+^CD25^+^ (**G**) cells of pLN from uninfected and IAV-infected mice (n= 6). **H**. Frequencies (left) and absolute cell counts (right) of CD11c^+^CII^+^ cDCs of pLN from uninfected and IAV-infected mice (n= 6). **I-K**. Frequencies (left) and absolute cell counts (right) of CD11c^+^CII^+^CD8^+^ cDC1 (**I**), CD11c^+^CII^+^ CD11b^+^ cDC2 (**J**), and CD11c^int^PDCA1^+^ pDCs (**K**) of pLN from uninfected and IAV-infected mice (n= 6). **L & M**. Surface expression levels (GMFI) of MHC-CI and MHC-CII (**L**), and CD80 and CD86 (**M**) in CD8^+^DCs (left), CD11b^+^DCs (middle) and CD11c^int^PDCA1^+^ pDCs from the pLN of uninfected and IAV-infected mice (n=6). Shown are data from mice after 10 days of IAV infection (**A-M**). All data represent mean ± SEM. Two-tailed student’s t tests were used to assess statistical significance (n. s. = not significant, *P < 0.05, **P<0.01, *** P< 0.001 & **** P< 0.0001).

Analysis of DC lineage revealed normal frequencies, but reduced absolute numbers, of CD11c^+^CII^+^ total cDCs (**Fig. 3H**); increased frequencies, but reduced absolute numbers, of CD8^+^CD11c^+^CII^+^ cDC1 (**Fig. 3I**); reduced frequencies and absolute numbers of CD11b^+^CD11c^+^CII^+^ cDC2 (**Fig. 3J**); and normal frequencies, but reduced absolute numbers of CD11c^low^PDCA1^+^ pDCs (**Fig. 3K**) within the lymph nodes of IAV-infected mice. Further analysis indicated elevated expression of MHC-Class I (**Fig. 3L; top**) and MHC-Class II (**Fig. 3L; bottom**) in cDC1, cDC2 and pDC subsets, increased expression of CD80 in cDC2 (**Fig. 3M; top**), and increased expression of CD86 in cDC2 and pDC subsets (**Fig. 3M; bottom**) in the lymph nodes of IAV-infected mice. These data specify that IAV infection leads to an imbalance of immune subsets of both lymphoid and myeloid lineages and upregulates activation markers in cDC and pDC subsets.

### IAV infection suppresses differentiation of immune cells in the thymus and BM

To assess if early T cell differentiation is altered in the thymus following IAV infection, we enumerated the frequencies and absolute numbers of T cell progenitors in the thymus. Our immunophenotyping studies indicated; increased frequencies, but normal absolute numbers, of CD4^+^CD8^-^ single positive (SP) 4 cells (**Fig. 4A, B**) and CD8^+^CD4^-^ SP8 cells (**Fig. 4A,C**); reduced frequencies and absolute numbers of CD4^+^CD8^+^ double positive (DP) cells (**Fig. 4 A, D**); and increased frequencies, but decreased absolute numbers, of CD4^-^CD8^-^ double negative (DN) cells (**Fig. 4 A, E**) within the thymus of IAV-infected mice. Further analysis of DN fractions (**Fig. 4F**) indicated increased frequencies, but normal absolute numbers, of CD44^+^CD25^-^ DN1 (**Fig. 4G**), normal frequencies and absolute numbers of CD44^+^CD25^+^ DN2 (**Fig. 4H**), and reduced frequencies and absolute numbers of CD44^-^CD25^+^ DN3 (**Fig. 4I**) and CD44^-^CD25^-^ DN4 (**Fig. 4J**) cells of IAV-infected mice.

**Figure 4.**
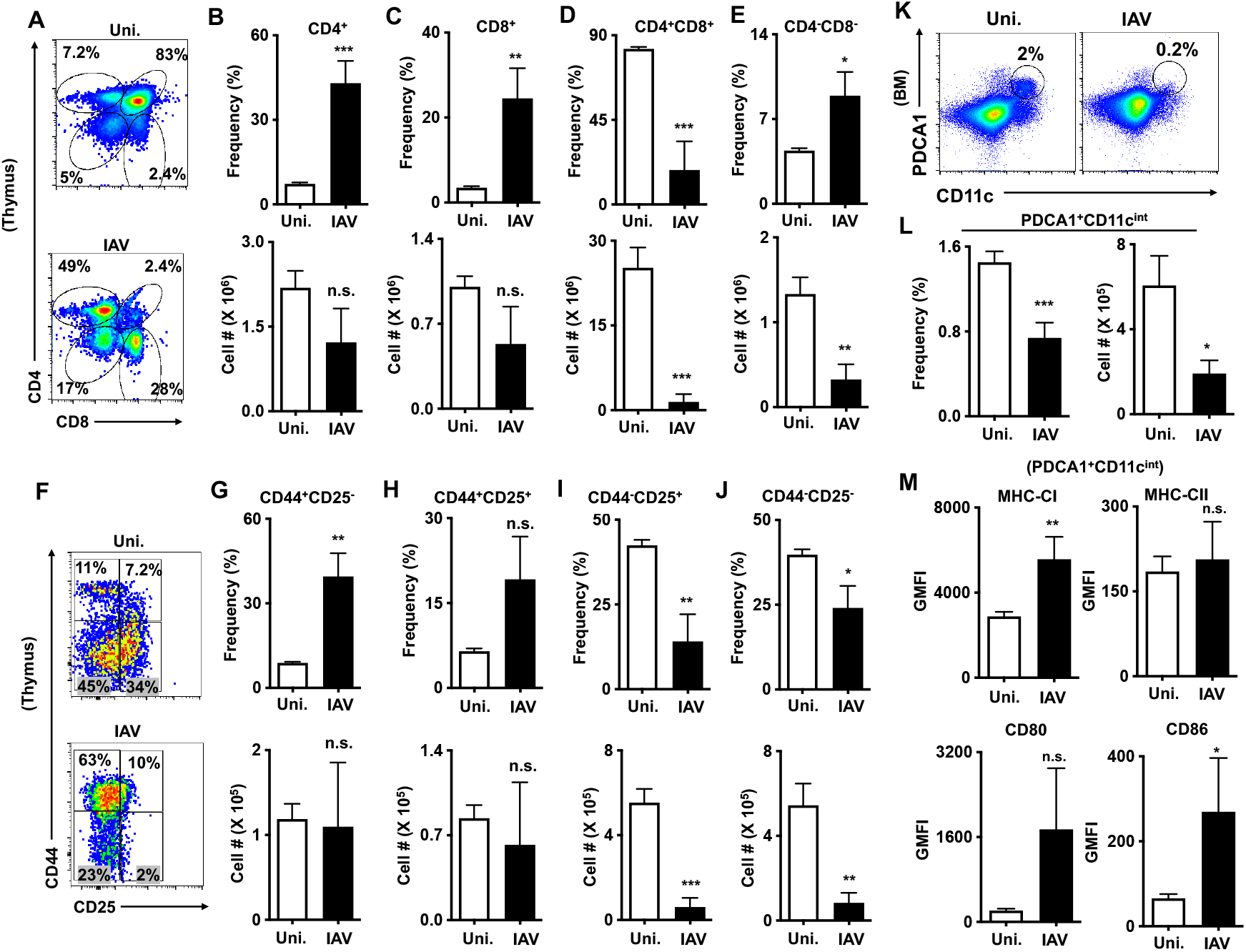
IAV infection leads to differentiation defects in the thymus and BM. **A**. FACS plots indicating the gating strategy of identifying CD4^+^CD8^-^ (SP4), CD4^-^CD8^+^ (SP8), CD4^+^CD8^+^ (DP) and CD4^-^CD8^-^ (DN) subsets from the thymus of uninfected and IAV-infected mice (n=6). **B-E**. Frequencies (left) and absolute cell counts (right) of SP4 (**B**), SP8 (**C**), DP (**D**) and DN (**E**) subsets from the thymus of uninfected and IAV-infected mice (n=6). **F**. FACS plots indicating the gating strategy of identifying CD44^+^CD25^-^ DN1, CD44^+^CD25^+^ DN2, CD44^-^ CD25^+^ DN3 and CD44^-^CD25^-^ DN4 subsets within CD4^-^CD8^-^ (DN) subset from the thymus of uninfected and IAV-infected mice (n=6). **G-J**. Frequencies (top) and absolute cell counts (bottom) of DN1 (**G**), DN2 (**H**), DN3 (**I**) and DN4 (**J**) subsets within CD4^-^CD8^-^ DN cells from the thymus of uninfected and IAV-infected mice (n=6). **K & L**. FACS plots indicating the gating strategy of identifying CD11c^int^PDCA1^+^ pDCs (**K**), and frequencies (left) and absolute cell counts (right) of pDCs (**L**) from the BM of uninfected and IAV-infected mice (n= 6). **M**. Surface expression levels (GMFI) of MHC-CI and MHC-CII, CD80 and CD86 in CD11c^int^PDCA1^+^ pDCs from the BM of uninfected and IAV-infected mice (n=6). Shown are data from mice after 10 days of IAV infection (**A-M**). All data represent mean ± SEM. Two-tailed student’s t tests were used to assess statistical significance (n. s. = not significant, *P < 0.05, **P<0.01, *** P< 0.001 & **** P< 0.0001).

Next, we studied if the differentiation program within the BM is intact during IAV infections. Our immunophenotyping analysis revealed normal frequencies and absolute numbers of CD19^+^ B cells (**Fig. S3A, B**). Further detailed analysis of B cell differentiation in the BM, as described earlier(32) indicated normal frequencies of Hardy’s- (CD43^+^B220^+^CD24^-^BP1^-^) Fraction A, (CD43^+^B220^+^CD24^+^BP1^-^) Fraction B, and (CD43^+^B220^+^CD24^+^BP1^+^) Fraction C (**Fig. S3C**) in the BM of IAV-infected mice. However, the frequencies of Hardy’s-(CD43^-^B220^+^IgM^-^IgD^-^) Fraction D and (CD43^-^B220^+^IgM^+^IgD^-^) Fraction E were reduced, and (CD43^-^B220^+^IgM^+^IgD^+^) Fraction F were increased in the BM of IAV-infected mice(**Fig. S3D**). Analysis of myeloid differentiation indicated normal frequencies and absolute numbers of Ly6G^+^CD11b^+^ granulocytes (**Fig. S3E**,**F**) and Ly6G^-^CD11b^+^ myelo/monocytes (**Fig. S3E, G**) in the BM of IAV-infected mice. Intriguingly, frequencies and absolute numbers of PDCA1^+^CD11c^int^ pDCs were reduced (**Fig. 4K, L**) and immunophenotyping studies revealed upregulation of MHC-Class I and CD86, while the expression levels of MHC-Class II and CD80 were normal, in pDCs (**Fig. 4M**) of the BM from IAV-infected mice. Overall, these data suggest that IAV infections impairs T cell differentiation program in the thymus and DC differentiation program in the BM.

### IAV infection alters hematopoietic stem and progenitor cell (HSPC) pool of the BM

To understand the consequences of IAV infection on early hematopoietic events, we studied the HSCs and lineage-committed progenitors within the BM. Our immunophenotyping studies indicated an increase in frequencies and absolute numbers of Lineage (Lin)^-^Sca1^+^c-Kit^+^ (LSK) HSPCs (**Fig. 5A,B**).

**Figure 5.**
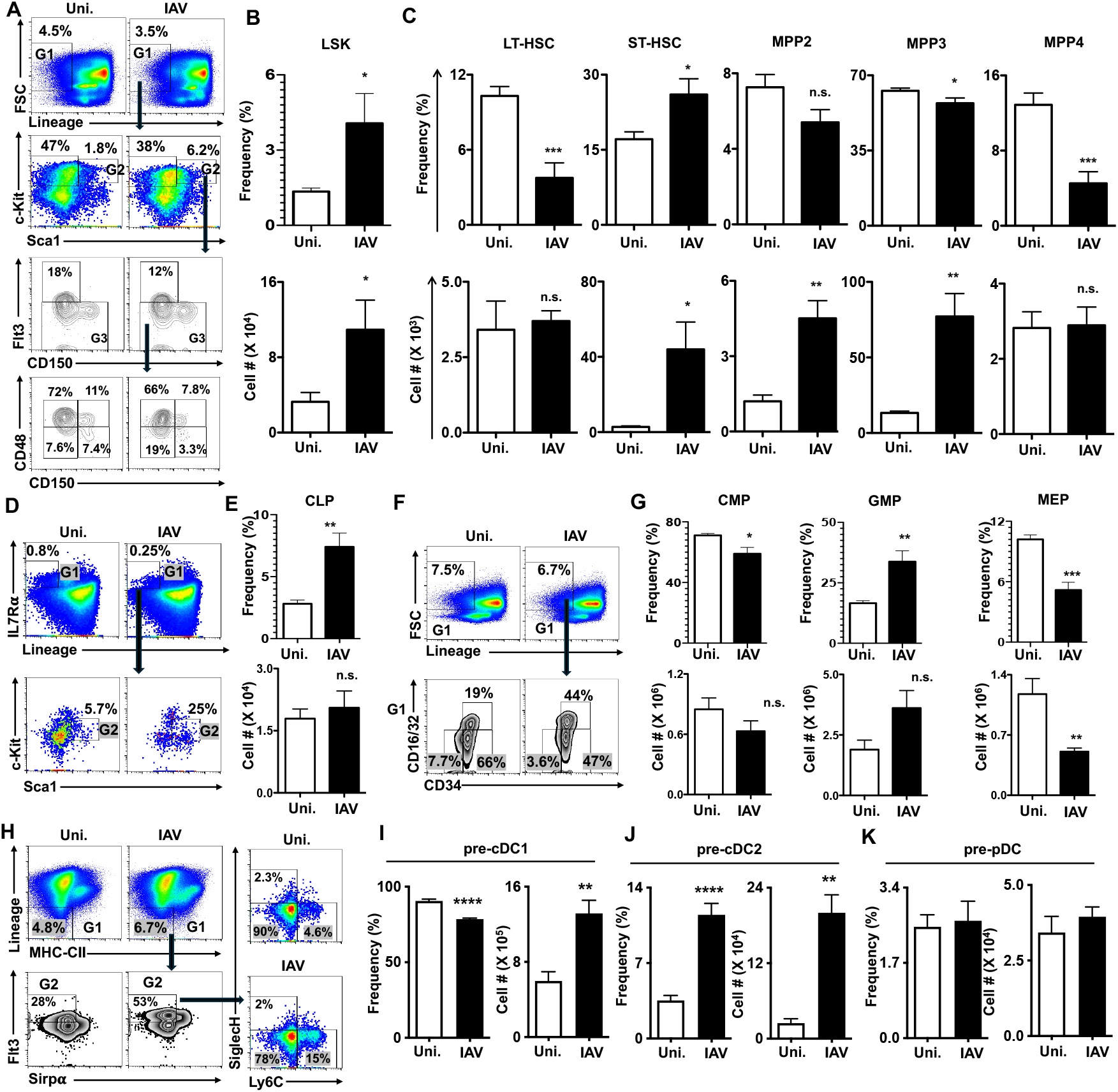
IAV infection results in altered HSPC subsets in the BM. **A**. FACS plots indicating the gating strategy of identifying HSPC subsets from the BM of uninfected and IAV-infected mice (n=6). **B**. Frequencies (top) and absolute cell counts (bottom) of LSK subset in the BM of uninfected and IAV-infected mice (n=6). **C**. Frequencies (top) and absolute cell counts (bottom) of LT-HSC, ST-HSC, MPP2, MPP3 and MPP4 subsets in the BM of uninfected and IAV-infected mice (n=6). **D**. FACS plots indicating the gating strategy of identifying CLP subset from the BM of uninfected and IAV-infected mice (n=6). **E**. Frequencies (top) and absolute cell counts (bottom) of CLP subset in the BM of uninfected and IAV-infected mice (n=6). **F**. FACS plots indicating the gating strategy of identifying CMP, GMP and MEP subsets from the BM of uninfected and IAV-infected mice (n=6). **G**. Frequencies (top) and absolute cell counts (bottom) of CMP, GMP and MEP subsets in the BM of uninfected and IAV-infected mice (n=6). **H**. FACS plots indicating the gating strategy of identifying pre-cDC1, pre-cDC2 and pre-pDC subsets from the BM of uninfected and IAV-infected mice (n=6). **I-K**. Frequencies (left) and absolute cell counts (right) of pre-cDC1 (**I**), pre-cDC2 (**J**) and pre-pDC (**K**) subsets in the BM of uninfected and IAV-infected mice (n=6). Shown are data from mice after 10 days of IAV infection (**A-K**). All data represent mean ± SEM. Two-tailed student’s t tests were used to assess statistical significance (n. s. = not significant, *P < 0.05, **P<0.01, *** P< 0.001 & **** P< 0.0001).

A detailed analysis(33) of LSK compartment indicated (**Fig. 5A,C**); reduced frequencies, but normal absolute numbers, of CD150^+^CD48^-^Flt3^-^LSK long-term HSCs (LT-HSCs); increased frequencies and absolute numbers of CD150^-^CD48^-^Flt3^-^LSK short-term HSCs (ST-HSCs); normal frequencies, but increased absolute numbers of CD150^+^CD48^+^Flt3^-^LSK multi-potent progenitors (MPP) 2; modestly reduced frequencies, but increased absolute numbers of CD150^-^CD48^+^Flt3^-^LSK MPP3; and reduced frequencies, but normal absolute numbers of Flt3^+^LSK MPP4, in the BM of IAV-infected mice. Next, we determined the numbers of lineage committed progenitors of the BM. Our immunophenotyping studies documented increased frequencies, but normal absolute numbers, of Lin^-^IL7Ra^+^Sca1^+^c-kit^int^ common lymphoid progenitors (CLPs) (**Fig. 5D, E**). Analysis of Lin^-^Sca1^-^c-kit^+^ (LK) myeloid progenitors indicated modestly reduced frequencies, but normal absolute numbers, of CD34^+^CD16/32^-^LK common myeloid progenitors (CMPs), increased frequencies, but normal absolute numbers, of CD34^+^CD16/32^+^LK granulocyte monocyte progenitors (GMPs), and reduced frequencies and absolute numbers of CD34^-^ CD16/32^-^LK megakaryocyte erythrocyte progenitors (MEPs) in the BM of IAV-infected mice (**Fig. 5F, G**). Finally, we determined the numbers of committed precursors (pre) of the cDC and pDC lineages(34). Our data indicated modestly reduced frequencies, but increased absolute numbers, of Lin^-^MHC-CII^-^ Flt3^+^Sirpa^-^ Ly6C^-^SiglecH^-^ pre-cDC1 (**Fig. 5H, I**), increased frequencies and absolute numbers of Lin^-^MHC-CII^-^ Flt3^+^Sirpa^-^ Ly6C^+^SiglecH^-^ pre-cDC2 (**Fig. 5H, J**), and normal frequencies and absolute numbers of MHC-CII^-^Flt3^+^Sirpa^-^Ly6C^-^SiglecH^+^ pre-pDC (**Fig. 5H, K**) in the BM of IAV-infected mice. These data indicated that IAV infection causes an imbalance in HSPC pool and alters the maintenance and early differentiation steps of DC subsets.

### IAV infection leads to increased systemic inflammation and exaggerated ROS levels in HSPCs

To understand the possible mechanisms through which IAV infection alters immune differentiation, we focused on the levels of inflammatory cytokines. For more than a decade, work of our lab(35-40), as well as of others(41-44), established that augmented systemic levels of inflammation cause major hematopoietic defects, such as loss of self-renewal, quiescence and impaired differentiation of HSCs, in the BM. Our ELISA studies demonstrated increased levels of Ifnα, Ifnβ, Ifnγ, Tnfα, IL18 and IL12, but normal levels of IL6 and IL1β, in the serum of IAV-infected mice (**Fig. 6A**). Next, we determined the local ‘pro-inflammatory niche’ of the lungs, spleen and BM through real-time (RT-) PCR studies. Our RT-PCR studies indicated increased levels of *Ifnα, Ifnβ* and *Ifnγ* mRNA and decreased levels of *Il6, Tnfα, Il1β, Il1β, Il18, Il33* and *Il10* mRNA in the lungs of IAV-infected mice (**Fig. 6B**). Intriguingly, analysis of spleen from IAV-infected mice revealed decreased levels of *Ifnβ, Ifnγ, Il6, Tnfα, Il1β* and *Il33* mRNA, and normal levels of *Ifnα, Il1β* and *Il10* mRNA (**Fig. 6C**). Analysis of BM from IAV-infected mice suggested increased levels of *Ifnα* and *Il18* mRNA, decreased levels of *Ifnβ, Ifnγ, Il1β, Il1β, Il33* and *Il10* mRNA, and normal levels of *Il6* and *Tnfα* mRNA (**Fig. 6D**). These data indicated that IAV infection results in augmented systemic levels of inflammatory cytokines and an imbalanced ‘pro-inflammatory cytokine milieu’ within the solid organs.

**Figure 6.**
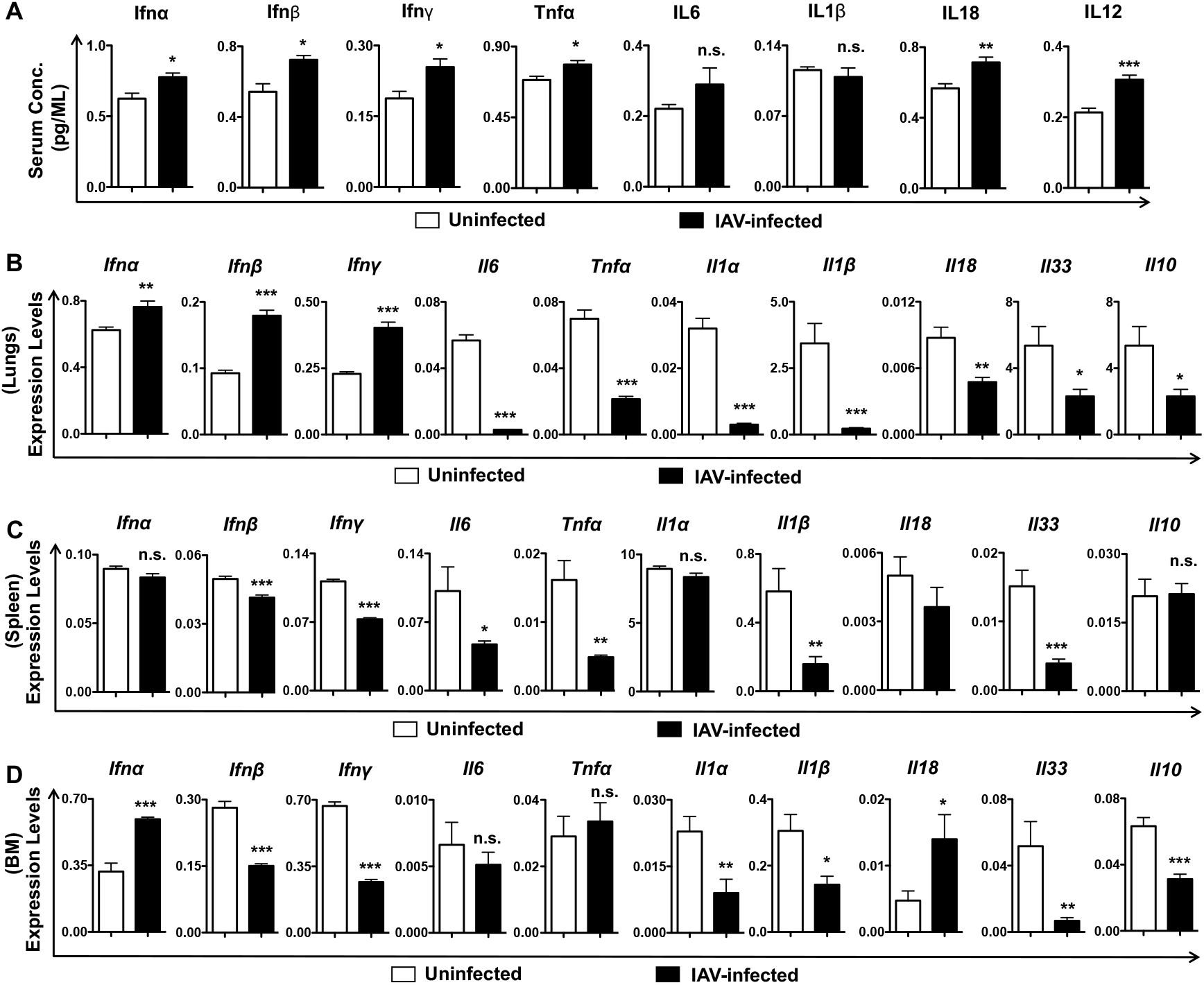
IAV infection causes elevated systemic inflammation and altered local inflammation in hematopoietic organs. **A**. ELISA data indicating circulating levels of pro-inflammatory cytokines in the blood serum of uninfected and IAV-infected mice (n=6). **B-D**. Real time PCR analysis documenting expression levels of *Ifnα, Ifnβ, Ifnγ, Il6, Tnfα, Il1β, Il1β, Il18, Il33* and *Il10* mRNA in the lungs (**B**), spleen (**C**) and BM (**D**) of uninfected and IAV-infected mice (n=6). Data were obtained using a pool of mRNA from 6 individual mice. Expression levels of target genes were normalized to *18s* RNA and *Actin* mRNA levels. Shown are data from mice after 10 days of IAV infection (**A-K**). All data represent mean ± SEM. Two-tailed student’s t tests were used to assess statistical significance (n. s. = not significant, *P < 0.05, **P<0.01, *** P< 0.001 & **** P< 0.0001).

To understand IAV-infected induced alterations at a molecular level, we determined the intracellular levels of ROS in HSPCs and differentiated immune cells. Indeed, ROS have been shown to play critical and decisive roles in the maintenance of HSPC pool and at the early differentiation stages of human and mouse DCs(45-48). Our flow cytometry studies indicated elevated levels of ROS in all stages of HSPCs, including Lin^-^, LK (Lin^-^c-Kit^+^), LSK, LT-HSCs, ST-HSCs, MPP2, MPP3 and MPP4 subsets of the BM from IAV-infected mice (**Fig. 7A**). On the other hand, analysis of differentiated cells indicated elevated levels of ROS in Ly6G^+^CD11b^+^ granulocytes, Ly6G^-^CD11b^+^ myelo/monocytes and CD19^+^ B cells, but normal levels of ROS in CD11c^+^ DCs, CD4^+^ T cells and CD8^+^ T cells, of the BM of IAV-infected mice (**Fig. 7B**). Analysis of immune subsets of the spleen indicated increased ROS levels in CD4^+^ T cells and CD8^+^ T cells, reduced ROS in CD11c^+^ DCs, and normal levels in Ly6G^+^CD11b^+^ granulocytes and Ly6G^-^ CD11b^+^ myelo/monocytes (**Fig. 7C**). Finally, we assessed if the increased ROS levels in HSPCs was due to increased active mitochondrial mass. Our flow analysis using Mito-Tracker-Green to selectively stain active mitochondria revealed increased mitochondrial mass in Lin-, LK, LSK, HSCs, MPP3 and MPP4 subsets of the BM from IAV-infected mice (**Fig. 7D & E**). In essence, these data suggest that IAV infection alters the mitochondrial mass and activity of early hematopoietic progenitors, including HSCs, that differentiate into mature cells of the myeloid and lymphoid lineages.

**Figure 7.**
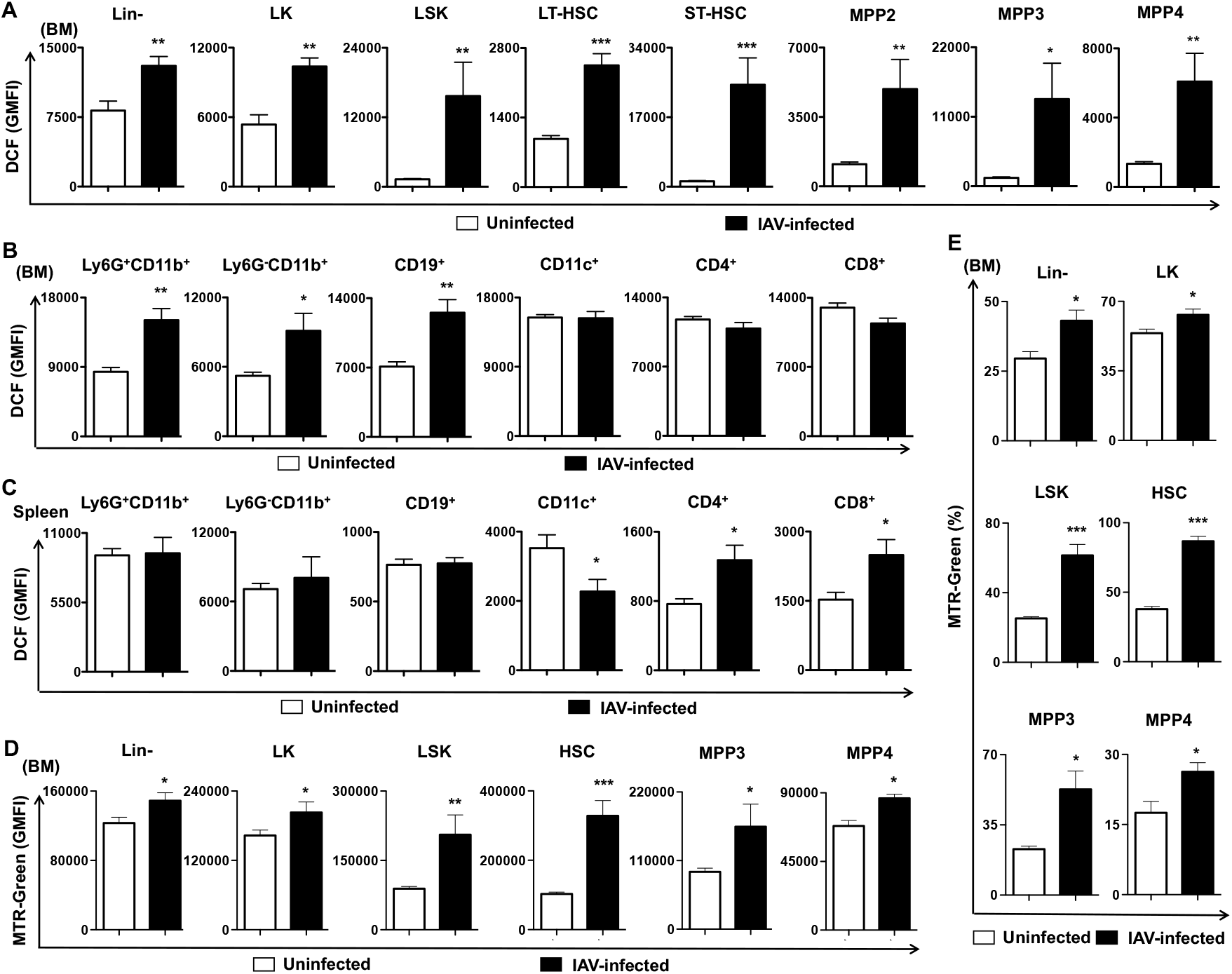
IAV infection results in augmented intracellular ROS levels and mitochondrial mass in HSPC subsets of the BM. **A**. Intracellular levels of ROS in lin-, LK, LSK, LT-HSC, ST-HSC, MPP2, MPP3 and MPP4 subsets in the BM of uninfected and IAV-infected mice (n=6). **B**. Intracellular levels of ROS in Ly6g^+^CD11b^+^, Ly6g^-^CD11b^+^, CD19^+^, CD11c^+^, CD4^+^ and CD8^+^ subsets in the BM of uninfected and IAV-infected mice (n=6). **C**. Intracellular levels of ROS in Ly6g^+^CD11b^+^, Ly6g^-^CD11b^+^, CD19^+^, CD11c^+^, CD4^+^ and CD8^+^ subsets in the spleen of uninfected and IAV-infected mice (n=6). **D**. Intracellular levels (GMFI) of Mitotracker green to detect mass of active mitochondria within lin-, LK, LSK, HSC, MPP3 and MPP4 subsets in the BM of uninfected and IAV-infected mice (n=6). **E**. Frequencies of Mitotracker green high lin-, LK, LSK, HSC, MPP3 and MPP4 subsets in the BM of uninfected and IAV-infected mice (n=6). Shown are data from mice after 10 days of IAV infection (**A-E**). All data represent mean ± SEM. Two-tailed student’s t tests were used to assess statistical significance (n. s. = not significant, *P < 0.05, **P<0.01, *** P< 0.001 & **** P< 0.0001).

## Discussion

Earlier studies demonstrated the involvement and importance of various immune cells, particularly DCs, in providing immunity and resistance against viral, particularly IAV, infections(1, 2, 22-26, 29). Among the three DC subsets; pDCs and two migratory DCs-CD103^+^ and CD11b^hi^ DCs(49) present in the naïve lungs, pulmonary CD103^+^ DCs of the lungs appear to play an indispensable role in providing immunity against IAV. CD103^+^ DCs cross-present antigens on MHC class I, whereas CD11b^hi^ DCs were responsible for antigen presentation on MHC class II(49). Indeed, CD103^+^ DCs have been demonstrated to control both priming of naive CD8^+^ T cells and regulate effector CD8^+^ T cell migration, survival, and memory responses during influenza infection(50-52). In vivo genetic ablation studies demonstrated that pulmonary CD103^+^ DCs are required for a cytotoxic T lymphocyte (CTL) response to viruses(50). Indeed, deficiency of CD103^+^ DCs caused a severe delay in the development of virus-specific CD8^+^ T cells and was correlated with increased clinical severity and a delayed viral clearance. Furthermore, CD103^+^ DCs of the lung capture and present apoptotic cell–associated antigen under homeostatic and inflammatory conditions(49). CD103^+^ DCs of the lungs play a key role in the transport of antigens from the lungs to the lymph nodes, and processing and presentation of viral antigens to CD8 T cells(53). In keeping with these findings, our data identified a severe reduction of CD103^+^ DCs in the lungs of mice that were infected with lethal dose of IAV infection. Intriguingly, we observed an increase of many other immune subsets including, CD11b^+^ DCs, pDCs, AVM, CD11b^+^ myelo/monocytes and CD8 T cells in the lungs after IAV infection. While increased infiltration and presence of immune cells in the lungs might compensate for the reduction of CD103^+^ DCs, it did not provide effective immunity against IAV and prevent lethality in mice. Even though pDCs were accumulated in the lungs during IAV infections, it appears to play a dispensable role in viral clearance or clinical severity(50), supporting the notion that CD103^+^ DCs might be the key mediator of anti-viral state in the lungs(51). In stark contrast, a few studies reported that both pDCs and cDCs play crucial roles in clearing IAV from the host(22, 54, 55). Therefore, more rigorous and in-depth studies focused on the precise involvement of specific DC subsets in providing immunity against IAV infections would be necessary to find novel treatment strategies.

IAV infection often predisposes individuals to secondary (bacterial) infections that lead to higher lethal outcomes(24). While the correlation between IAV and bacterial infections has been well known and recognized for the past several decades(24, 56), cellular and molecular mechanisms leading to the suppression of innate immune functions following IAV infection remain largely unknown(57-60). Emerging studies establish that IAV infection also leads to alterations in the distribution of immune cells in the peripheral lymphoid and non-lymphoid organs(15-17). However, the pathophysiological impact of IAV infection (and associated inflammation) in the bone marrow, particularly on HSC physiology, remains elusive. To date, only one study by Rommel et al. demonstrated(61) that IAV infection causes hyper-proliferation and preferential differentiation of HSCs into megakaryocytes (platelets) through IL1R-dependent mechanisms. However, the study of Rommel et al. did not document and discuss global alterations of the innate and adaptive immune cells following IAV infection, which might be critical in understanding IAV-induced immune defects and increased susceptibilities to secondary infections. To this end, studies described here were designed to identify alterations in the differentiation and/or maintenance of major immune subsets in lymphoid organs during lethal IAV infections. Our studies indicated that IAV infection leads to major alterations in immune cell differentiation and maintenance in the peripheral lymphoid organs. Strikingly, our data identified pancytopenia, as evidenced by major reduction in peripheral blood counts and severely diminished cellularity of the spleen, thymus and lymph nodes. Our evaluation of immune cell distribution in primary and secondary lymphoid organs indicated a severe imbalance in proportion and reduction in absolute numbers of both innate and adaptive immune cells. Based on our data, we hypothesize that pancytopenia of hematopoietic and immune cells might be one of the major contributing factors of lethality following IAV infections. Accelerated loss of peripheral immune cells during IAV infection might be explained by increased migration and recruitment to the lungs, and augmented apoptosis of immune cells. Indeed, apoptosis-induced during IAV infection play a critical role in cell death and tissue damage(62). On the other hand, our studies highlighted that early hematopoietic differentiation in the BM and T cell differentiation in the thymus were diminished during IAV infection. Intriguingly, our analysis on hematopoietic progenitors and lineage restricted precursor suggested that differentiation of specific immune subsets, such as DCs and T cells, is diminished/altered at crucial stages of their ontogeny. Therefore, we hypothesize that impaired differentiation of hematopoietic/immune cells might also contribute to ‘IAV-induced pancytopenia’ and immune defects. Further understanding on precise mechanisms leading to pancytopenia might be important in the treatment and management of this disease. Our mechanistic data identified elevated levels of systemic inflammation, altered levels of local inflammation in the BM, augmented levels of intracellular ROS and increased mass of active mitochondria in HSCs, hematopoietic progenitors of the myeloid, lymphoid and erythroid lineages. It was interesting to observe rather reduced levels of intracellular ROS in splenic DCs. ROS are key signaling molecules that play a key role in pathogenesis and there exists an interesting crosstalk between inflammation and ROS. Proinflammatory cytokines, such as Ifnα, IFNγ and Tnfα, can trigger and utilize ROS to influence immune cells under both physiological and inflammatory conditions(63, 64). For more than a decade, research(41-44), including our own(35-40), has established that augmented systemic inflammation impairs HSC pool and their differentiation into innate and adaptive immune cells. Furthermore, increased intracellular ROS levels compromise HSC maintenance and functions(65). Based on these findings, we believe that IAV-mediated inflammatory cascades might cause HSC defects and interfere with their differentiation into functionally competent immune cells. Nevertheless, we cannot exclude the possible involvement of additional mechanisms, such as direct sensing of IAV-infection by HSCs, through which IAV affects hematopoiesis and immune differentiation. Further in-depth mechanistic studies would be necessary and beneficial in understanding and targeting IAV-induced pathologies, especially involving the lung-BM axis.

## Footnotes

This work was supported by the National Heart, Lung, and Blood Institute research grant HL132194, and National Institute of Allergy and Infectious Diseases research grants-AI174952 and AI183982.

P.S., G.S. P.K. and N.S. performed research. C.V.R. conceptualized and designed research, acquired funding, performed research, collected data, analyzed data, interpreted data and wrote manuscript. All authors have read and agreed to the published version of the manuscript.

## References

1. Cole, S. L., J. Dunning, W. L. Kok, K. H. Benam, A. Benlahrech, E. Repapi, F. O. Martinez, L. Drumright, T. J. Powell, M. Bennett, R. Elderfield, C. Thomas, M. investigators, T. Dong, J. McCauley, F. Y. Liew, S. Taylor, M. Zambon, W. Barclay, V. Cerundolo, P. J. Openshaw, A. J. McMichael, and L. P. Ho. 2017. M1-like monocytes are a major immunological determinant of severity in previously healthy adults with life-threatening influenza. JCI Insight 2: e91868.

2. Oshansky, C. M., A. J. Gartland, S. S. Wong, T. Jeevan, D. Wang, P. L. Roddam, M. A. Caniza, T. Hertz, J. P. Devincenzo, R. J. Webby, and P. G. Thomas. 2014. Mucosal immune responses predict clinical outcomes during influenza infection independently of age and viral load. Am J Respir Crit Care Med 189: 449–462.

3. Guo, X. J., and P. G. Thomas. 2017. New fronts emerge in the influenza cytokine storm. Semin Immunopathol 39: 541–550.

4. Damjanovic, D., C. L. Small, M. Jeyanathan, S. McCormick, and Z. Xing. 2012. Immunopathology in influenza virus infection: uncoupling the friend from foe. Clin Immunol 144: 57–69.

5. Liu, Q., Y. H. Zhou, and Z. Q. Yang. 2016. The cytokine storm of severe influenza and development of immunomodulatory therapy. Cell Mol Immunol 13: 3–10.

6. Tscherne, D. M., and A. Garcia-Sastre. 2011. Virulence determinants of pandemic influenza viruses. J Clin Invest 121: 6–13.

7. Walsh, K. B., J. R. Teijaro, P. R. Wilker, A. Jatzek, D. M. Fremgen, S. C. Das, T. Watanabe, M. Hatta, K. Shinya, M. Suresh, Y. Kawaoka, H. Rosen, and M. B. Oldstone. 2011. Suppression of cytokine storm with a sphingosine analog provides protection against pathogenic influenza virus. Proc Natl Acad Sci U S A 108: 12018–12023.

8. D’Elia, R. V., K. Harrison, P. C. Oyston, R. A. Lukaszewski, and G. C. Clark. 2013. Targeting the “cytokine storm” for therapeutic benefit. Clin Vaccine Immunol 20: 319–327.

9. Teijaro, J. R. 2015. The role of cytokine responses during influenza virus pathogenesis and potential therapeutic options. Curr Top Microbiol Immunol 386: 3–22.

10. Tisoncik, J. R., M. J. Korth, C. P. Simmons, J. Farrar, T. R. Martin, and M. G. Katze. 2012. Into the eye of the cytokine storm. Microbiol Mol Biol Rev 76: 16–32.

11. Davison, A. M., D. Thomson, and J. S. Robson. 1973. Intravascular coagulation complicating influenza A virus infection. Br Med J 1: 654–655.

12. Short, K. R., E. Kroeze, R. A. M. Fouchier, and T. Kuiken. 2014. Pathogenesis of influenza-induced acute respiratory distress syndrome. Lancet Infect Dis 14: 57–69.

13. Watanabe, T. 2013. Renal complications of seasonal and pandemic influenza A virus infections. Eur J Pediatr 172: 15–22.

14. La Gruta, N. L., K. Kedzierska, J. Stambas, and P. C. Doherty. 2007. A question of self-preservation: immunopathology in influenza virus infection. Immunol Cell Biol 85: 85–92.

15. Cheung, C. Y., L. L. Poon, A. S. Lau, W. Luk, Y. L. Lau, K. F. Shortridge, S. Gordon, Y. Guan, and J. S. Peiris. 2002. Induction of proinflammatory cytokines in human macrophages by influenza A (H5N1) viruses: a mechanism for the unusual severity of human disease? Lancet 360: 1831–1837.

16. Sakabe, S., K. Iwatsuki-Horimoto, R. Takano, C. A. Nidom, M. T. Q. Le, T. Nagamura-Inoue, T. Horimoto, N. Yamashita, and Y. Kawaoka. 2011. Cytokine production by primary human macrophages infected with highly pathogenic H5N1 or pandemic H1N1 2009 influenza viruses. J Gen Virol 92: 1428–1434.

17. Wang, J., M. P. Nikrad, E. A. Travanty, B. Zhou, T. Phang, B. Gao, T. Alford, Y. Ito, P. Nahreini, K. Hartshorn, D. Wentworth, C. A. Dinarello, and R. J. Mason. 2012. Innate immune response of human alveolar macrophages during influenza A infection. PLoS One 7: e29879.

18. Doherty, P. C., D. J. Topham, R. A. Tripp, R. D. Cardin, J. W. Brooks, and P. G. Stevenson. 1997. Effector CD4+ and CD8+ T-cell mechanisms in the control of respiratory virus infections. Immunol Rev 159: 105–117.

19. Bender, B. S., and P. A. Small, Jr. 1992. Influenza: pathogenesis and host defense. Semin Respir Infect 7: 38–45.

20. Kwissa, M., H. I. Nakaya, N. Onlamoon, J. Wrammert, F. Villinger, G. C. Perng, S. Yoksan, K. Pattanapanyasat, K. Chokephaibulkit, R. Ahmed, and B. Pulendran. 2014. Dengue virus infection induces expansion of a CD14(+)CD16(+) monocyte population that stimulates plasmablast differentiation. Cell Host Microbe 16: 115–127.

21. Sporri, R., and C. Reis e Sousa. 2005. Inflammatory mediators are insufficient for full dendritic cell activation and promote expansion of CD4+ T cell populations lacking helper function. Nat Immunol 6: 163–170.

22. Hemann, E. A., L. E. Sjaastad, R. A. Langlois, and K. L. Legge. 2015. Plasmacytoid Dendritic Cells Require Direct Infection To Sustain the Pulmonary Influenza A Virus-Specific CD8 T Cell Response. J Virol 90: 2830–2837.

23. Kawasaki, T., M. Ikegawa, and T. Kawai. 2022. Antigen Presentation in the Lung. Front Immunol 13: 860915.

24. Smed-Sorensen, A., C. Chalouni, B. Chatterjee, L. Cohn, P. Blattmann, N. Nakamura, L. Delamarre, and I. Mellman. 2012. Influenza A virus infection of human primary dendritic cells impairs their ability to cross-present antigen to CD8 T cells. PLoS Pathog 8: e1002572.

25. Vangeti, S., S. Falck-Jones, M. Yu, B. Osterberg, S. Liu, M. Asghar, K. Sonden, C. Paterson, P. Whitley, J. Albert, N. Johansson, A. Farnert, and A. Smed-Sorensen. 2023. Human influenza virus infection elicits distinct patterns of monocyte and dendritic cell mobilization in blood and the nasopharynx. Elife 12.

26. Wakim, L. M., and M. J. Bevan. 2011. Cross-dressed dendritic cells drive memory CD8+ T-cell activation after viral infection. Nature 471: 629–632.

27. Dunning, J., S. Blankley, L. T. Hoang, M. Cox, C. M. Graham, P. L. James, C. I. Bloom, D. Chaussabel, J. Banchereau, S. J. Brett, M. F. Moffatt, A. O’Garra, P. J. M. Openshaw, and M. Investigators. 2018. Progression of whole-blood transcriptional signatures from interferon-induced to neutrophil-associated patterns in severe influenza. Nat Immunol 19: 625–635.

28. Gill, M. A., K. Long, T. Kwon, L. Muniz, A. Mejias, J. Connolly, L. Roy, J. Banchereau, and O. Ramilo. 2008. Differential recruitment of dendritic cells and monocytes to respiratory mucosal sites in children with influenza virus or respiratory syncytial virus infection. J Infect Dis 198: 1667–1676.

29. Gill, M. A., A. K. Palucka, T. Barton, F. Ghaffar, H. Jafri, J. Banchereau, and O. Ramilo. 2005. Mobilization of plasmacytoid and myeloid dendritic cells to mucosal sites in children with respiratory syncytial virus and other viral respiratory infections. J Infect Dis 191: 1105–1115.

30. Chen, W. H., F. R. Toapanta, K. A. Shirey, L. Zhang, A. Giannelou, C. Page, M. B. Frieman, S. N. Vogel, and A. S. Cross. 2012. Potential role for alternatively activated macrophages in the secondary bacterial infection during recovery from influenza. Immunol Lett 141: 227–234.

31. Lewis, K. L., M. L. Caton, M. Bogunovic, M. Greter, L. T. Grajkowska, D. Ng, A. Klinakis, I. F. Charo, S. Jung, J. L. Gommerman, Ivanov, II, K. Liu, M. Merad, and B. Reizis. 2011. Notch2 receptor signaling controls functional differentiation of dendritic cells in the spleen and intestine. Immunity 35: 780–791.

32. Hardy, R. R., and K. Hayakawa. 2001. B cell development pathways. Annu Rev Immunol 19: 595–621.

33. Pietras, E. M., D. Reynaud, Y. A. Kang, D. Carlin, F. J. Calero-Nieto, A. D. Leavitt, J. M. Stuart, B. Gottgens, and E. Passegue. 2015. Functionally Distinct Subsets of Lineage-Biased Multipotent Progenitors Control Blood Production in Normal and Regenerative Conditions. Cell Stem Cell 17: 35–46.

34. Schlitzer, A., V. Sivakamasundari, J. Chen, H. R. Sumatoh, J. Schreuder, J. Lum, B. Malleret, S. Zhang, A. Larbi, F. Zolezzi, L. Renia, M. Poidinger, S. Naik, E. W. Newell, P. Robson, and F. Ginhoux. 2015. Identification of cDC1- and cDC2-committed DC progenitors reveals early lineage priming at the common DC progenitor stage in the bone marrow. Nat Immunol 16: 718–728.

35. Nakagawa, M. M., H. Chen, and C. V. Rathinam. 2018. Constitutive Activation of NF-kappaB Pathway in Hematopoietic Stem Cells Causes Loss of Quiescence and Deregulated Transcription Factor Networks. Front Cell Dev Biol 6: 143.

36. Nakagawa, M. M., H. Davis, and C. V. Rathinam. 2018. A20 deficiency in multipotent progenitors perturbs quiescence of hematopoietic stem cells. Stem Cell Res 33: 199–205.

37. Nakagawa, M. M., and C. V. Rathinam. 2018. Constitutive Activation of the Canonical NF-kappaB Pathway Leads to Bone Marrow Failure and Induction of Erythroid Signature in Hematopoietic Stem Cells. Cell Rep 25: 2094–2109 e2094.

38. Nakagawa, M. M., and C. V. Rathinam. 2019. A20 deficiency in hematopoietic stem cells causes lymphopenia and myeloproliferation due to elevated Interferon-gamma signals. Sci Rep 9: 12658.

39. Nakagawa, M. M., K. Thummar, J. Mandelbaum, L. Pasqualucci, and C. V. Rathinam. 2015. Lack of the ubiquitin-editing enzyme A20 results in loss of hematopoietic stem cell quiescence. J Exp Med 212: 203–216.

40. Rathinam, C. 2015. The ‘inflammatory’ control of hematopoietic stem cells. Oncotarget 6: 19938–19939.

41. de Bruin, A. M., S. F. Libregts, M. Valkhof, L. Boon, I. P. Touw, and M. A. Nolte. 2012. IFNgamma induces monopoiesis and inhibits neutrophil development during inflammation. Blood 119: 1543–1554.

42. de Bruin, A. M., C. Voermans, and M. A. Nolte. 2014. Impact of interferon-gamma on hematopoiesis. Blood 124: 2479–2486.

43. King, K. Y., and M. A. Goodell. 2011. Inflammatory modulation of HSCs: viewing the HSC as a foundation for the immune response. Nat Rev Immunol 11: 685–692.

44. Mirantes, C., E. Passegue, and E. M. Pietras. 2014. Pro-inflammatory cytokines: emerging players regulating HSC function in normal and diseased hematopoiesis. Exp Cell Res 329: 248–254.

45. Del Prete, A., P. Zaccagnino, M. Di Paola, M. Saltarella, C. Oliveros Celis, B. Nico, G. Santoro, and M. Lorusso. 2008. Role of mitochondria and reactive oxygen species in dendritic cell differentiation and functions. Free Radic Biol Med 44: 1443–1451.

46. Sattler, M., T. Winkler, S. Verma, C. H. Byrne, G. Shrikhande, R. Salgia, and J. D. Griffin. 1999. Hematopoietic growth factors signal through the formation of reactive oxygen species. Blood 93: 2928–2935.

47. Sheng, K. C., G. A. Pietersz, C. K. Tang, P. A. Ramsland, and V. Apostolopoulos. 2010. Reactive oxygen species level defines two functionally distinctive stages of inflammatory dendritic cell development from mouse bone marrow. J Immunol 184: 2863–2872.

48. Sinha, A., A. Singh, V. Satchidanandam, and K. Natarajan. 2006. Impaired generation of reactive oxygen species during differentiation of dendritic cells (DCs) by Mycobacterium tuberculosis secretory antigen (MTSA) and subsequent activation of MTSA-DCs by mycobacteria results in increased intracellular survival. J Immunol 177: 468–478.

49. Desch, A. N., G. J. Randolph, K. Murphy, E. L. Gautier, R. M. Kedl, M. H. Lahoud, I. Caminschi, K. Shortman, P. M. Henson, and C. V. Jakubzick. 2011. CD103+ pulmonary dendritic cells preferentially acquire and present apoptotic cell-associated antigen. J Exp Med 208: 1789–1797.

50. GeurtsvanKessel, C. H., M. A. Willart, L. S. van Rijt, F. Muskens, M. Kool, C. Baas, K. Thielemans, C. Bennett, B. E. Clausen, H. C. Hoogsteden, A. D. Osterhaus, G. F. Rimmelzwaan, and B. N. Lambrecht. 2008. Clearance of influenza virus from the lung depends on migratory langerin+CD11b-but not plasmacytoid dendritic cells. J Exp Med 205: 1621–1634.

51. Helft, J., B. Manicassamy, P. Guermonprez, D. Hashimoto, A. Silvin, J. Agudo, B. D. Brown, M. Schmolke, J. C. Miller, M. Leboeuf, K. M. Murphy, A. Garcia-Sastre, and M. Merad. 2012. Cross-presenting CD103+ dendritic cells are protected from influenza virus infection. J Clin Invest 122: 4037–4047.

52. Ng, S. L., Y. J. Teo, Y. A. Setiagani, K. Karjalainen, and C. Ruedl. 2018. Type 1 Conventional CD103(+) Dendritic Cells Control Effector CD8(+) T Cell Migration, Survival, and Memory Responses During Influenza Infection. Front Immunol 9: 3043.

53. Ho, A. W., N. Prabhu, R. J. Betts, M. Q. Ge, X. Dai, P. E. Hutchinson, F. C. Lew, K. L. Wong, B. J. Hanson, P. A. Macary, and D. M. Kemeny. 2011. Lung CD103+ dendritic cells efficiently transport influenza virus to the lymph node and load viral antigen onto MHC class I for presentation to CD8 T cells. J Immunol 187: 6011–6021.

54. McGill, J., and K. L. Legge. 2009. Cutting edge: contribution of lung-resident T cell proliferation to the overall magnitude of the antigen-specific CD8 T cell response in the lungs following murine influenza virus infection. J Immunol 183: 4177–4181.

55. McGill, J., N. Van Rooijen, and K. L. Legge. 2008. Protective influenza-specific CD8 T cell responses require interactions with dendritic cells in the lungs. J Exp Med 205: 1635–1646.

56. Beadling, C., and M. K. Slifka. 2004. How do viral infections predispose patients to bacterial infections? Curr Opin Infect Dis 17: 185–191.

57. Didierlaurent, A., J. Goulding, S. Patel, R. Snelgrove, L. Low, M. Bebien, T. Lawrence, L. S. van Rijt, B. N. Lambrecht, J. C. Sirard, and T. Hussell. 2008. Sustained desensitization to bacterial Toll-like receptor ligands after resolution of respiratory influenza infection. J Exp Med 205: 323–329.

58. Jamieson, A. M., S. Yu, C. H. Annicelli, and R. Medzhitov. 2010. Influenza virus-induced glucocorticoids compromise innate host defense against a secondary bacterial infection. Cell Host Microbe 7: 103–114.

59. Shahangian, A., E. K. Chow, X. Tian, J. R. Kang, A. Ghaffari, S. Y. Liu, J. A. Belperio, G. Cheng, and J. C. Deng. 2009. Type I IFNs mediate development of postinfluenza bacterial pneumonia in mice. J Clin Invest 119: 1910–1920.

60. van der Sluijs, K. F., L. J. van Elden, M. Nijhuis, R. Schuurman, J. M. Pater, S. Florquin, M. Goldman, H. M. Jansen, R. Lutter, and T. van der Poll. 2004. IL-10 is an important mediator of the enhanced susceptibility to pneumococcal pneumonia after influenza infection. J Immunol 172: 7603–7609.

61. Rommel, M. G. E., L. Walz, F. Fotopoulou, S. Kohlscheen, F. Schenk, C. Miskey, L. Botezatu, Y. Krebs, I. M. Voelker, K. Wittwer, T. Holland-Letz, Z. Ivics, V. von Messling, M. A. G. Essers, M.D. Milsom, C. K. Pfaller, and U. Modlich. 2022. Influenza A virus infection instructs hematopoiesis to megakaryocyte-lineage output. Cell Rep 41: 111447.

62. Tran, A. T., J. P. Cortens, Q. Du, J. A. Wilkins, and K. M. Coombs. 2013. Influenza virus induces apoptosis via BAD-mediated mitochondrial dysregulation. J Virol 87: 1049–1060.

63. Abad, C., I. Pinal-Fernandez, C. Guillou, G. Bourdenet, L. Drouot, P. Cosette, M. Giannini, L. Debrut, L. Jean, S. Bernard, D. Genty, R. Zoubairi, I. Remy-Jouet, B. Geny, C. Boitard, A. Mammen, A. Meyer, and O. Boyer. 2024. IFNgamma causes mitochondrial dysfunction and oxidative stress in myositis. Nat Commun 15: 5403.

64. Blaser, H., C. Dostert, T. W. Mak, and D. Brenner. 2016. TNF and ROS Crosstalk in Inflammation. Trends Cell Biol 26: 249–261.

65. Ludin, A., S. Gur-Cohen, K. Golan, K. B. Kaufmann, T. Itkin, C. Medaglia, X. J. Lu, G. Ledergor, O. Kollet, and T. Lapidot. 2014. Reactive oxygen species regulate hematopoietic stem cell self-renewal, migration and development, as well as their bone marrow microenvironment. Antioxid Redox Signal 21: 1605–1619.

