## Supplemental Files for "Influenza virus infection in the lungs leads to pancytopenia and defective immune cell differentiation program in the thymus and bone marrow"

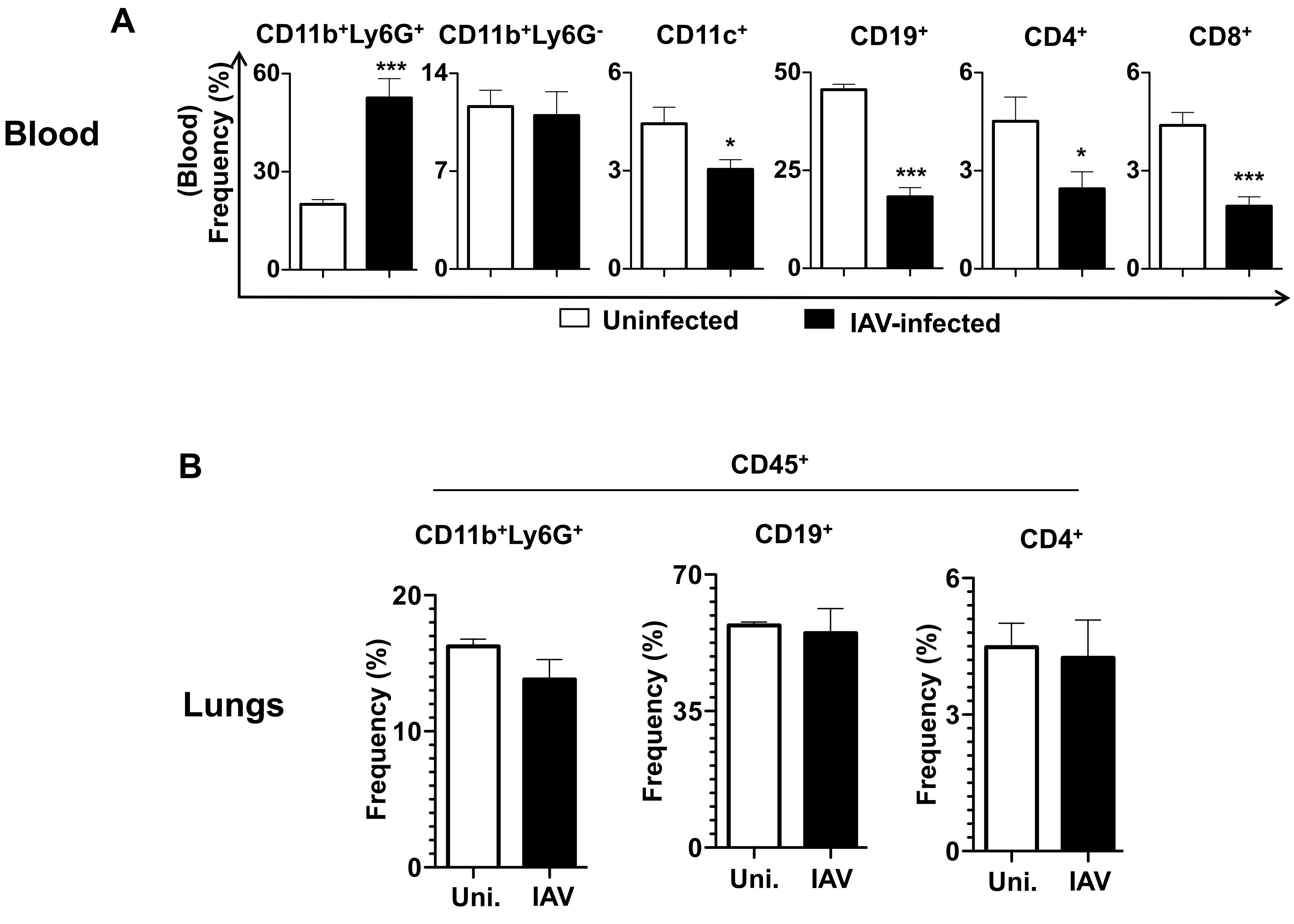
A
Blood
CD11b+Ly6G+
CD11b+Ly6G-
CD11c+
CD19+
CD4+
CD8+
(Blood) Frequency (%)
Uninfected
IAV-infected
B
CD45+
Lungs
CD11b+Ly6G+
CD19+
CD4+
Frequency (%)
Uni.
IAV

### Figure S1.

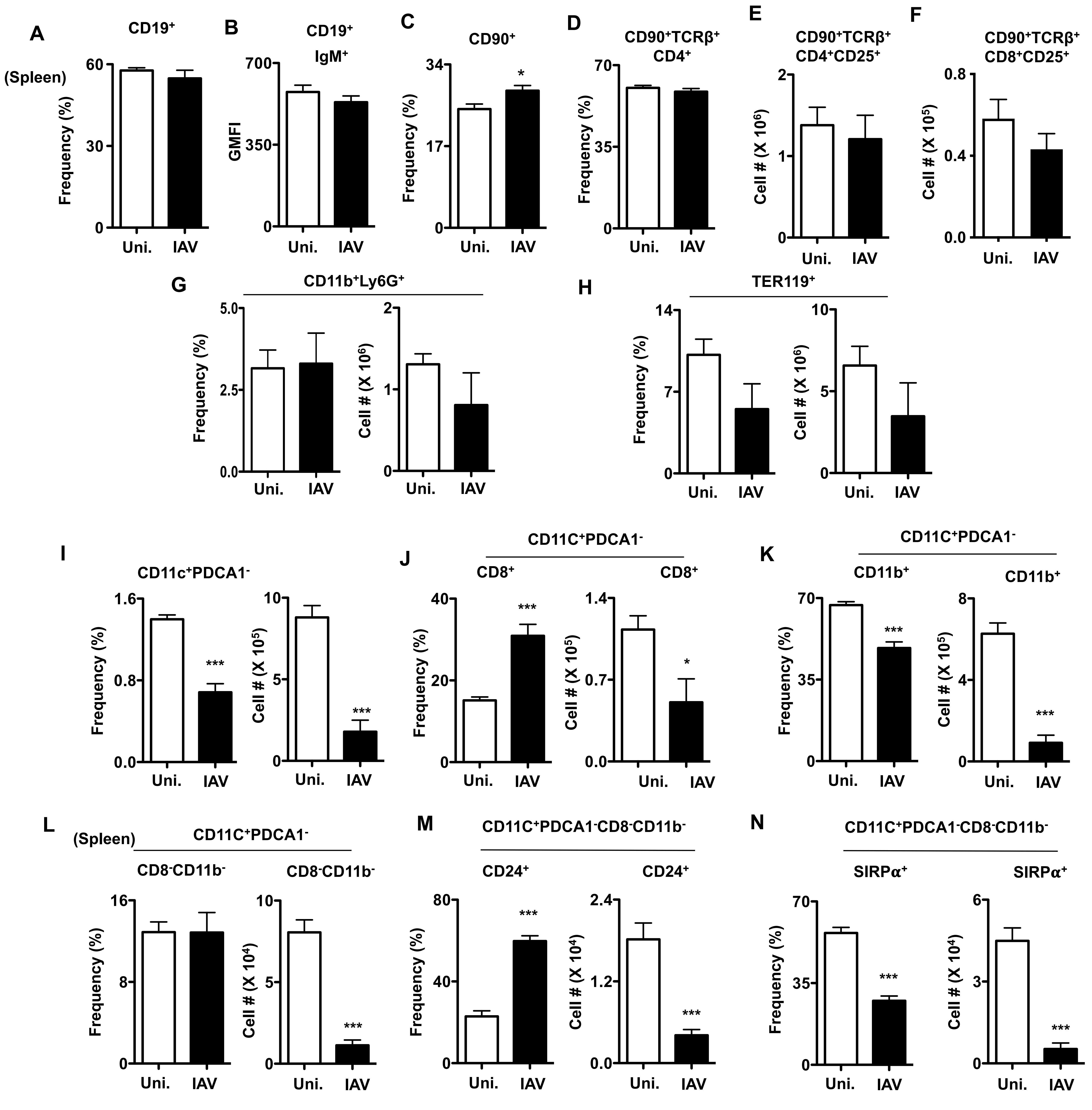
A
CD19+
(Spleen)
Frequency (%)
Uni.
IAV
B
CD19+ IgM+
GMFI
C
CD90+
D
CD90+TCRβ+ CD4+
E
CD90+TCRβ+ CD4+CD25+
Cell # (X 10^6)
F
CD90+TCRβ+ CD8+CD25+
Cell # (X 10^5)
G
CD11b+Ly6G+
H
TER119+
I
CD11c+PDCA1-
J
CD11C+PDCA1-
CD8+
K
CD11C+PDCA1-
CD11b+
L
(Spleen)
CD11C+PDCA1-
CD8-CD11b-
Cell # (X 10^4)
M
CD11C+PDCA1-CD8-CD11b-
CD24+
N
CD11C+PDCA1-CD8-CD11b-
SIRPα+

### Figure S2.

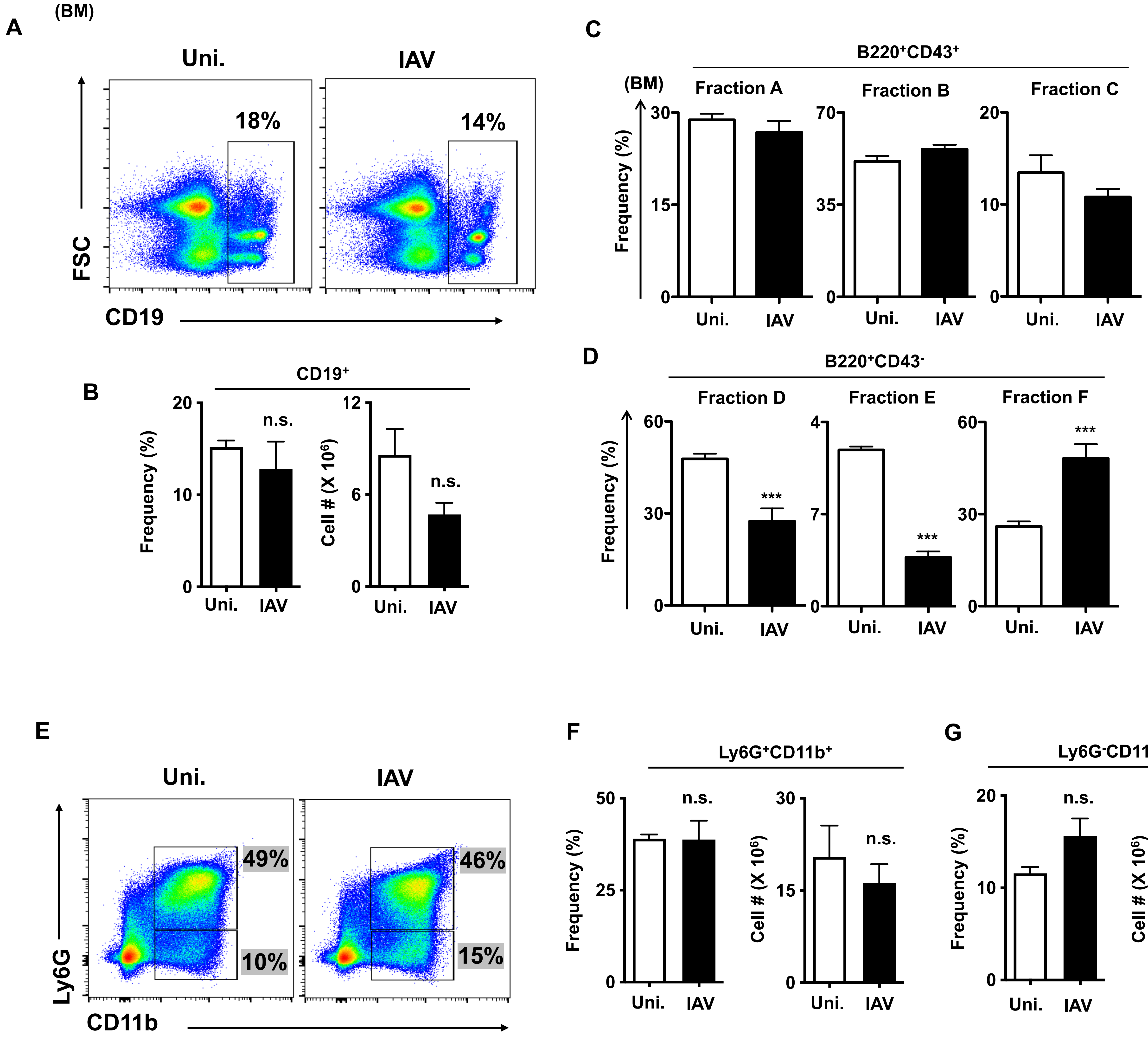
A
(BM)
Uni.
IAV
18%
14%
FSC
CD19
B
CD19⁺
Frequency (%)
20
10
0
n.s.
Uni.
IAV
Cell # (X 10⁶)
12
6
0
n.s.
Uni.
IAV
C
B220⁺CD43⁺
(BM)
Fraction A
Fraction B
Fraction C
Frequency (%)
30
15
0
70
35
0
20
10
0
Uni.
IAV
D
B220⁺CD43⁻
Fraction D
Fraction E
Fraction F
Frequency (%)
60
30
0
4
7
0
60
30
0
***
***
***
Uni.
IAV
E
Uni.
IAV
49%
10%
46%
15%
Ly6G
CD11b
F
Ly6G⁺CD11b⁺
Frequency (%)
50
25
0
n.s.
Cell # (X 10⁶)
30
15
0
n.s.
Uni.
IAV
G
Ly6G⁻CD11b⁺
Frequency (%)
20
10
0
n.s.
Cell # (X 10⁶)
8
4
0
n.s.
Uni.
IAV

Figure S3.

#### Supplemental Figure legends:

**Figure S1.**

**A.** Frequencies of Ly6g$^+$CD11b$^+$, Ly6g$^-$CD11b$^+$, CD11c$^+$, CD19$^+$, CD4$^+$ and CD8$^+$ subsets in the peripheral blood of uninfected and IAV-infected mice (n=6).

**B.** Frequencies of Ly6g$^+$CD11b$^+$, CD19$^+$ and CD4$^+$ subsets within CD45$^+$ cells of the lungs of uninfected and IAV-infected mice (n=6).

**Figure S2.**

**A.** Frequencies of CD19$^+$ subset in the spleen of uninfected and IAV-infected mice (n=6).

**B.** Surface expression levels (GMFI) of IgM in the spleen of uninfected and IAV-infected mice (n=6).

**C.** Frequencies of CD90$^+$ subset in the spleen of uninfected and IAV-infected mice (n=6).

**D.** Frequencies of CD90$^+$TCRb$^+$CD4$^+$ subset in the spleen of uninfected and IAV-infected mice (n=6).

**E.** Absolute cell counts of CD90$^+$TCRb$^+$CD4$^+$CD25$^+$ subset in the spleen of uninfected and IAV-infected mice (n=6).

**F.** Absolute cell counts of CD90$^+$TCRb$^+$CD8$^+$CD25$^+$ subset in the spleen of uninfected and IAV-infected mice (n=6).

**G.** Frequencies (left) absolute cell counts (right) of Ly6g$^+$CD11b$^+$ subset in the spleen of uninfected and IAV-infected mice (n=6).

**H.** Frequencies (left) absolute cell counts (right) of Ter119$^+$ subset in the spleen of uninfected and IAV-infected mice (n=6).

**I.** Frequencies (left) absolute cell counts (right) of CD11C$^{int}$PDCA1$^-$ subset in the spleen of uninfected and IAV-infected mice (n=6).

**J.** Frequencies (left) absolute cell counts (right) of CD8$^+$ cDC1 subset within CD11C$^{int}$PDCA1$^-$ fraction in the spleen of uninfected and IAV-infected mice (n=6).

**K.** Frequencies (left) absolute cell counts (right) of CD11b$^+$ cDC2 subset within CD11C$^{int}$PDCA1$^-$ fraction in the spleen of uninfected and IAV-infected mice (n=6).

**L.** Frequencies (left) absolute cell counts (right) of CD8$^-$CD11b$^-$ cells within CD11C$^{int}$PDCA1$^-$ fraction in the spleen of uninfected and IAV-infected mice (n=6).

**M.** Frequencies (left) absolute cell counts (right) of CD24$^+$Sirp$\alpha^-$ cells within CD8$^-$CD11b$^-$ CD11C$^{int}$PDCA1$^-$ fraction in the spleen of uninfected and IAV-infected mice (n=6).

**N.** Frequencies (left) absolute cell counts (right) of Sirp$\alpha^+$CD24$^-$ cells within CD8$^-$CD11b$^-$ CD11C$^{int}$PDCA1$^-$ fraction in the spleen of uninfected and IAV-infected mice (n=6).

Shown are data from mice after 10 days of IAV infection (A-N). All data represent mean ± SEM. Two-tailed student's t tests were used to assess statistical significance (n. s. = not significant, $*P < 0.05$, $**P<0.01$, $***P< 0.001$ & $****P< 0.0001$).

**Figure S3.**

**A.** FACS plots indicating the gating strategy of identifying CD19$^+$ B cells from the BM of uninfected and IAV-infected mice (n=6).

**B.** Frequencies (left) absolute cell counts (right) of CD19$^+$ B cells in the BM of uninfected and IAV-infected mice (n=6).

**C.** Frequencies of CD24$^-$BP1$^-$ Fraction-A, CD24$^+$BP1$^-$ Fraction-B, CD24$^+$BP1$^+$ Fraction-C within the CD43$^+$B220$^+$ fraction of the BM of uninfected and IAV-infected mice (n=6).

**D.** Frequencies of IgM$^-$IgD$^-$ Fraction-D, IgM$^+$IgD$^-$ Fraction-E, IgM$^+$IgD$^+$ Fraction-F within the CD43$^-$B220$^+$ fraction of the BM of uninfected and IAV-infected mice (n=6).

**E.** FACS plots indicating the gating strategy of identifying Ly6g$^+$CD11b$^+$ and Ly6g$^-$CD11b$^+$ cells from the BM of uninfected and IAV-infected mice (n=6).

**F.** Frequencies (left) absolute cell counts (right) of Ly6g$^{+}$CD11b$^{+}$ subset in the BM of uninfected and IAV-infected mice (n=6).

**G.** Frequencies (left) absolute cell counts (right) of Ly6g$^{-}$CD11b$^{+}$ subset in the BM of uninfected and IAV-infected mice (n=6).

Shown are data from mice after 10 days of IAV infection (A-G). All data represent mean ± SEM. Two-tailed student's t tests were used to assess statistical significance (n. s. = not significant, *P < 0.05, **P<0.01, *** P< 0.001 & **** P< 0.0001).
